# Conservation and divergence in cortical cellular organization between human and mouse revealed by single-cell transcriptome imaging

**DOI:** 10.1101/2021.11.01.466826

**Authors:** Rongxin Fang, Chenglong Xia, Meng Zhang, Jiang He, Jennie Close, Brian Long, Jeremy Miller, Ed Lein, Xiaowei Zhuang

## Abstract

The human cerebral cortex has tremendous cellular diversity and complex cellular organization that are essential for brain function. How different types of cells are organized and interact with each other in the human cortex, and how cellular organizations and interaction patterns vary across species are, however, unclear. Here, we performed spatially resolved single-cell expression profiling of 4,000 genes in human middle and superior temporal gyrus using multiplexed error-robust fluorescence *in situ* hybridization (MERFISH). We identified >100 neuronal and non-neuronal cell populations with distinct transcriptional signatures, generated a molecularly defined and spatially resolved cell atlas of these brain regions, and analyzed cell-cell interactions in a cell-type-specific manner. Comparison with the mouse cortex showed conservation in the laminar organization of cells and substantial divergence in cell-cell interactions between human and mouse. Notably, our data revealed a drastic increase in interactions between neurons and non-neuronal cells in the human cortex, uncovered human-specific cell-cell interaction patterns, and identified potential ligand-receptor basis of microglia-neuron interactions.

## Main Text

The human cerebral cortex comprises billions of cells of distinct types and their spatial organizations and interactions play critical roles in shaping and maintaining various brain functions (*1, 2*). For instance, interactions between neuronal and non-neuronal cells are essential for axonal conduction, synaptic transmission, and information processing, and are therefore required for normal functioning of the nervous system during development and throughout adult life (*3, 4*). Disruption of such cell-cell interactions has recently been shown to contribute to various neurological disorders, such as autism (*5*), schizophrenia (*6*), and Alzheimer’s diseases (*7*). Despite their importance, we only have a limited understanding of the organizations and interactions of different types of cells in the human cortex, and how they vary across species and relate to brain diseases.

Recent single-cell RNA sequencing analysis has revealed a high diversity of transcriptionally distinct cell populations in the human MTG (*8*), a model system for studying human cortex because fresh surgical tissues are often available for this brain region. Combination of single-cell RNA sequencing with microdissection (*8*), and more recently an *in situ* sequencing study targeting 120 genes (*9*), have revealed the laminar organization of these transcriptionally defined neuronal cell types, in particular the excitatory neuronal cell types, in the human MTG. However, a systematic understanding of the spatial relationship and cell-cell interactions among this high diversity of cell types is still lacking. Single-cell transcriptomics and epigenomics analyses have also provided rich insights into the evolution of cellular diversity and molecular signatures of cell types in mouse, marmoset monkey and human cortex (*8, 10, 11*), but how the spatial relationship and interactions between different cell types vary across species remains largely unclear.

Recent advances in spatially resolved single-cell transcriptomics have enabled *in situ* gene expression profiling of individual cells and spatial mapping of cell types in complex tissues (*12*). Among these, multiplexed error-robust fluorescence *in situ* hybridization (MERFISH) is a single-cell transcriptome-imaging method that allows simultaneous imaging of thousands of genes in individual cells (*13, 14*). MERFISH has been applied to map the spatial organizations of cell types in mouse brain tissues (*15, 16*). Here, we demonstrated MERFISH imaging of human brain tissues and profiled the expression of 4,000 genes in the human cortex using this approach. The MERFISH images enabled *in situ* identification of >100 neuronal and non-neuronal cell populations the human MTG and superior temporal gyrus (STG), comprehensive mapping of the spatial organization of these cell populations, and systematic characterization of proximity-based cell-cell interactions in a cell-type specific manner. Furthermore, cross-species comparison based on the MERFISH data revealed both common features and striking differences in the composition, spatial organizations, and interaction patterns of cell types in the human and mouse cortex.

We performed MERFISH measurements of the human MTG and STG from both fresh-frozen neurosurgical and postmortem brain samples, targeting 4,000 genes (**Fig. 1A**). These genes included 764 marker genes differentially expressed in the cell clusters derived from single-nucleus SMART-seq data of the human MTG (*8*) and 3,236 additional genes expressed in this region largely randomly selected to increase the gene coverage (**Materials and Methods**). This allowed us to include potential marker genes not identified in the SMART-seq data (for example, due to the depletion of non-neuronal cells from the SMART-seq samples), as well as functionally important genes such as ligand and receptor pairs. Because human neurons are known to have high autofluorescence due to the aging pigments such as lipofuscin aggregates (*17*), detection of individual RNA molecules is challenging for human brain tissues. To overcome this challenge, we illuminated the samples with multi-band light emitting diode arrays (*18*) to eliminate the autofluorescence background prior to MERFISH probe staining and imaging. We then used expansion microscopy (*19*) to reduce the molecular crowding effect associated with imaging such a large number of genes (*14, 20*).

**Fig. 1.**
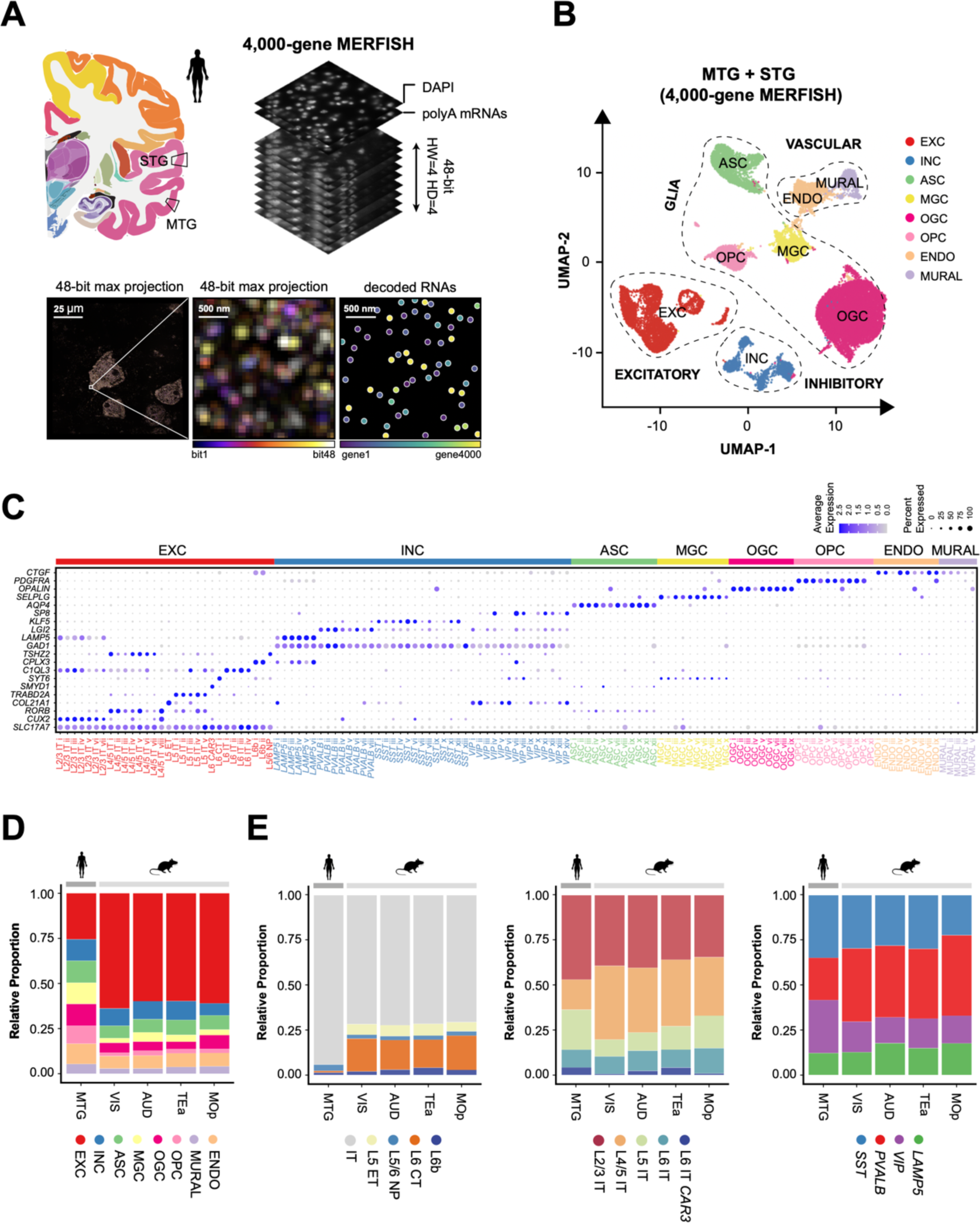
Spatially resolved single-cell transcriptome profiling of the human and mouse cortex by MERFISH. (**A**) (Top left) Schematic of the middle temporal gyrus (MTG) and superior temporal gyrus (STG) region of the human cortex. Black boxes indicate the area dissected for MERFISH imaging. (Top right) Schematic of the MERFISH measurements. 4,000 genes were imaged by MERFISH using a 48-bit hamming distance (HW) of 4 and hamming code (HD) of 4 code. Total polyadenylated RNA (polyA) and DAPI imaging was used to identify cell boundaries. (Bottom left) A 4000-gene MERFISH image of a human MTG sample in a single z-plane of a single field-of-view, with the maximum intensity across the 48 bits displayed in each pixel. (Bottom middle) A zoomed-in image of the boxed region in the left image. (Bottom right) RNA molecules decoded in the middle image. The scale bar indicates the real size of the sample prior to the sample expansion. (**B**) UMAP visualization of excitatory neurons, inhibitory neurons and major subclasses of non-neuronal cells identified in the human MTG and STG using the 4,000-gene MERFISH datasets. EXC: excitatory neurons; INC: inhibitory neurons; ASC: astrocytes; MGC: microglial cells; OGC: oligodendrocytes; OPC: oligodendrocyte progenitor cells; ENDO: endothelial cells; and MURAL: mural cells. (**C**) Subdivisions of the major subclasses of cells into clusters and the expression of a selective subset of marker genes in these clusters displayed by dot plot. The size of each dot corresponds to the percentage of cells expressing the gene in each cluster and the color indicates the average normalized expression level. (**D**) MERFISH-derived proportions of the excitatory neurons, inhibitory neurons, and major subclasses of non-neuronal cells in human MTG and four mouse cortical areas including primary motor cortex (MOp), visual cortex (VIS), auditory cortex (AUD) and temporal association area (TEa). (**E**) Proportion of subclasses of excitatory neurons (left), subclasses of IT neurons (middle), and subclasses of inhibitory neurons (right) in human MTG and the four mouse cortical areas.

Individual RNA molecules were clearly detected in these samples and decoding of the MERFISH images allowed us to determine the identity of these RNA molecules (**Fig. 1A**). We segmented individual cells in the tissue sections based on the 4′,6-diamidino-2-phenylindole (DAPI) and total polyadenylated RNA staining using a deep learning segmentation algorithm (*21*) (**Fig. S1**) and assigned RNA molecules to segmented cells to determine the single-cell expression profiles *in situ*. MERFISH expression data showed excellent reproducibility between replicates (**Fig. S2A**), and good correlation with bulk RNA sequencing data (**Fig. S2B**). We found that the median transcript counts per cell in the postmortem samples was ∼53% of that of the fresh-frozen neurosurgical samples (**Fig. S2C**), possibly due to RNA degradation during the processing of the postmortem brain. Despite the reduction in total RNA count, the expression levels of individual genes showed high correlation between the postmortem and fresh samples (**Fig. S2C**), allowing us to combine results from both types of samples. To test whether the molecular crowding associated with imaging 4,000 genes caused a substantial reduction in the detection efficiency, we performed MERFISH imaging on 250 genes chosen from the 764 marker genes derived from SMART-seq data. The detection efficiency of the 4000-gene measurements was on average 57% of that determined by the 250-gene measurements (**Fig. S3**). Moreover, the expression levels of individual genes were highly correlated between the 4000-gene and 250-gene measurements (**Fig. S3**).

Next, we identified transcriptionally distinct cell populations using the single-cell expression profiles derived from the 4,000-gene MERFISH data. We identified excitatory (EXC) and inhibitory (INC) neurons, as well as major subclasses of non-neuronal cells including microglia (MGC), astrocytes (ASC), oligodendrocytes (OGC), oligodendrocyte progenitor cells (OPC), endothelial cells (ENDO) and mural cells (MURAL), as characterized by the marker genes previously identified by SMART-seq (*8*) (**Fig. 1B** **and Fig. S4**). We then performed separate clustering analyses of the inhibitory and excitatory neurons using the MERFISH data (**Fig. S5, A and B**) and the SMART-seq data (**Fig. S5, C and D**) independently. The clusters determined by MERFISH showed good correspondence to those independently determined by SMART-seq (**Fig. S5, E and F**). To combine information from both types of measurements, we performed integrative analysis of the 4,000-gene MERFISH data and the SMART-seq data to jointly define clusters within the excitatory and inhibitory neuronal populations (**Fig. S6, A and B**). This integrative analysis classified inhibitory neurons into four subclasses, denoted by their markers *SST*, *VIP*, *PVALB* and *LAMP5*, with each subclass further divided into multiple clusters (**Fig. 1C**). Similarly, the excitatory neurons were classified into nine subclasses (L2/3 IT, L4/5 IT, L5 IT, L6 IT, L6 IT *CAR3*, L5 ET, L5/6 NP, L6 CT, L6b) based on their known marker genes, with most subclasses also sub-divided into multiple clusters (**Fig. 1C**). Because non-neuronal cells were depleted from the samples used for SMART-seq analysis (*8*), we identified clusters within individual subclasses of the non-neuronal cells based on the 4,000-gene MERFISH data alone (**Fig. 1C** **and Fig. S6C**). Using this approach, we identified a total of 127 transcriptionally distinct cell populations in human MTG and STG, including 30 excitatory, 41 inhibitory, 56 non-neuronal clusters (**Fig. 1C**), revealing not only a high diversity of neurons, but also a remarkably high diversity of non-neuronal cells in the human cortex. In addition, to include our 250-gene MERFISH datasets for the downstream analysis, we performed supervised classification to predict the cluster labels for cells measured in the 250-gene experiments based on the annotation of the 4,000-gene MERFISH datasets.

Next, we used our MERFISH data to provide a quantitative analysis of the cell composition in the human MTG. Because different surgical samples often include different amount of white matter, which is predominantly made of non-neuronal cells, we excluded the white matter from our cell composition analysis to avoid arbitrariness in the non-neuronal cell proportion. Overall, the human MTG was composed of 23% excitatory, 11% inhibitory, and 66% non-neuronal cells (**Fig. 1D**). The excitatory neurons were predominantly IT neurons (∼94%), with only a small fraction of non-IT neurons (1.2% L6 CT, 0.2% L5 ET, 3.5% L5/6 NP and 1.5% L6b cells) (**Fig. 1E, left**). The IT neurons were sub-divided into 47% L2/3 IT, 17% L4/5 IT, 22% L5 IT, 10% L6 IT and 4% L6 IT *CAR3* (**Fig. 1E, middle**). The inhibitory neurons were composed of 12% *LAMP5*, 23% *PVALB*, 35% *SST*, 30% *VIP* cells (**Fig. 1E, right**).

Next, we compared the cell composition between human and mouse cortices. Because the human MTG does not have a direct homologous area in the mouse brain, we considered two MERFISH datasets covering several regions of the mouse cortex. The first one is our recently reported 258-gene MERFISH dataset covering the primary motor cortex (MOp) and adjacent areas (*16*). In addition, we performed MERFISH experiments on a more posterior part of the mouse cortex, including the visual cortex (VIS), auditory cortex (AUD), temporal association area (TEa), and other cortical areas (**Fig. S7, A-B**). For cell classification of these newly imaged mouse cortical regions, we performed integrative analysis of the MERFISH and SMART-seq data (*8*) (**Fig. S7, C-F**) in the same manner as we analyzed the human MTG and STG datasets. We observed similar cell compositions across the different mouse cortical regions imaged, including MOp, VIS, AUD, and TEa (**Fig. 1, D and E**).

The human MTG, however, exhibited substantial differences in the cell composition compared to these mouse cortical regions. Notably, we observed a much lower proportion of excitatory neurons and a substantially higher proportion of the glial cells (including astrocytes, oligodendrocytes, OPCs and microglia) in the human cortex as compared to the mouse cortex (**Fig. 1D**). The glia-to-neuron ratio was 1.5 in the human cortex, consistent with results from other cell counting methods (*22, 23*), but this ratio was much higher than the 0.3 value observed for the mouse cortex (**Fig. 1D**). The excitatory-to-inhibitory neuron ratio was 2:1 in human, in line with recent independent measurements (*9, 11*) and substantially lower than the 6:1 ratio observed in mouse (**Fig. 1D**). Among the excitatory neurons, the non-IT neuron proportion dropped drastically to only 6% of excitatory neurons in the human cortex, from 29% in the mouse cortex (**Fig. 1E, left**). These results were consistent with recent observations that L5 ET and L6 CT are much less abundant in the primates compared to mouse (*11*). The dominance of IT neurons and the low proportions of ET and CT neurons in the human cortex suggest a dramatic increase in the fraction of neurons specialized for intra-cortical communication compared to those specialized for cortical-subcortical connections in human. For inhibitory neurons, we observed a substantial decrease in the proportion of *PVALB* neurons and a substantial increase in the proportion of *VIP* neurons in human compared to mouse (**Fig. 1E, right**). *VIP* interneurons regulate feedback inhibition of excitatory neurons through suppression of other interneurons such as *SST* and *PVALB* (*24*), thereby modulating network dynamics based on the behavioral state (*25, 26*). The observed increase of *VIP* interneurons in the human cortex thus provides a potential mechanism for human’s enhanced ability in state-dependent sensory processing and learning-related neuronal dynamics.

*In situ* identification of cell types by MERFISH allowed us to map the spatial organization of these cells. Among the excitatory neurons, we observed a laminar organization of the IT neurons in the human MTG, with individual IT clusters often sub-dividing a cortical layer into finer laminar structures and neighboring IT clusters showing overlapping spatial distributions (**Fig. 2, A left and B top**). The non-IT neurons, including L5 ET, L5/6 NP, L6 CT and L6b, were populated mostly in the deep layers (L5 and L6), as expected (**Fig. 2, A left and B top**). Among inhibitory neurons, *VIP* and *LAMP5* were enriched in upper layers (L1-L3) while *PVALB* and *SST* were broadly distributed from L2 to L6 (**Fig. 2, A middle and B middle**) (*8, 9*). Notably, at the cluster level, inhibitory neurons also adopted a pronounced laminar organization, with many inhibitory clusters being primarily restricted to one cortical layer, or even a sub-portion of a layer (**Fig. 2B** **middle**). These spatial organizations of excitatory and inhibitory neurons were overall similar to those observed in the mouse cortex (**Fig. S8B**) (*16*).

**Fig. 2.**
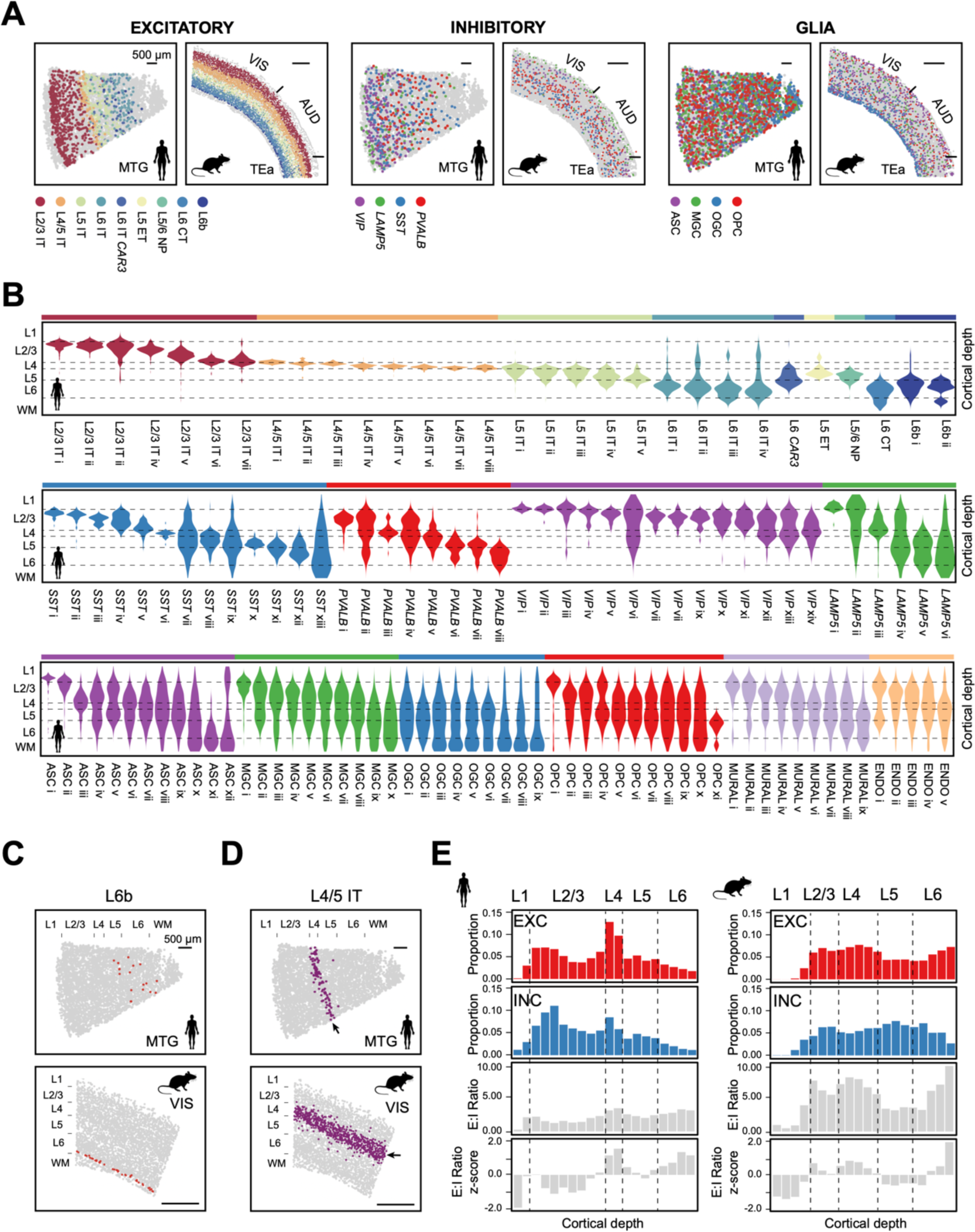
Spatial organization of cell types in the human and mouse cortex. (**A**) Spatial maps of the subclasses of excitatory neurons (left two panels), inhibitory neurons (middle two panels) and glial cells (right two panels) determined by MERFISH in a human MTG slice and a mouse slice that contained the VIS, AUD and TEa. Indicated subclasses of cells are shown in colors and other cells are shown in grey. (**B**) Cortical-depth distribution of excitatory (top), inhibitory (middle) and non-neuronal (bottom) clusters in the human MTG shown as violin plots. The cortical depths of cells were normalized in each slice and the dashed grey lines mark the approximate layer boundaries. WM: white matter. (**C-D**) Spatial maps of L6b (**C**) and L4/5 IT (**D**) neurons in a human MTG slice (top) and a VIS-containing region in a mouse slice (bottom). (**E**) Normalized cortical-depth distributions of excitatory (top row) and inhibitory (second row) neurons, E:I ratio (third row), and E:I ratio z-score (bottom row) along the cortical depth axis in human (left) and mouse cortex (right). E:I ratio was determined as the ratio between the number of excitatory and inhibitory neurons in each cortical depth bin.

Despite this overall conservation for laminar organization of neurons in human and mouse cortices, we also found marked differences for some cell types. For instance, the excitatory neuronal type L6b was broadly dispersed in L6 and extending into L5 and the white matter in human MTG, whereas in mouse cortex L6b formed a very thin layer at the bottom border of L6 (**Fig. 2C**). This is in line with the previous characterizations of the subplate neurons in the human cortex (*27*) and the L6b marker gene *CTGF* in the mouse and human cortex (*28*). Although the proportion of L4/5 IT neurons decrease in human compared to that in mouse (**Fig. 1E, middle**), surprisingly, we found that the L4/5 IT neurons formed a very dense and thin layer in the human MTG, giving rise to a substantially higher density of excitatory neurons in L4 than in other cortical layers in human, whereas the density of excitatory neurons in the mouse cortex was more uniform across L2/3 to L6 (**Fig. 2, D and E top**). As L4 is known to vary between different cortical regions, whether this difference is region- or species-specific remains an open question although the several mouse cortical regions that we examined exhibited a similar density profile. In addition, we also found that the cortical-depth dependence of the excitatory-to-inhibitory neuron ratio (E:I) was substantially different between human and mouse (**Fig. 2E**).

Notably, non-neuronal cell types also exhibited laminar organization in the human cortex. Oligodendrocytes were enriched in the white matter and depleted in the upper layers (L1-L3) (**Fig. S9**) (*9*). Although astrocytes, microglia, OPCs, endothelial cells and mural cells appeared dispersed across all cortical layers at the subclass level (**Fig. S9**), interestingly, these cell types exhibited a laminar organization at the cluster level (**Fig. 2B** **bottom**). Taking astrocytes as an example, the ASC i cluster was highly enriched in L1 and near the boundary between L1 and L2/3, likely representing interlaminar astrocytes (*8*), ASC ii was enriched in lower L1 and upper L2/3, ASC iii was enriched in lower L2/3 and L4, ASC iv – ix were dispersed across L2/3-L6, and ASC x – xii were enriched in L6 and the white matter (**Fig. 2B** **bottom**). Similar to astrocytes, nearly all non-neuronal subclasses exhibited a gradually evolving cell composition across the cortical depth (**Fig. 2B** **bottom**).

Importantly, high-resolution measurements of the spatial relationship between cells by MERFISH allowed us to predict cell-cell interactions arising from somatic contact or paracrine signaling, which can be inferred based on contact or proximity between cells that occurred with higher frequency than by random chance (*29, 30*). Our MERFISH images showed frequent somatic contacts among neurons, among non-neuronal cells, as well as between neurons and non-neuronal cells (**Fig. 3A**). We examined whether these potential cell-cell interactions were cell-type specific. To this end, we only considered cell types at the subclass level and determined the frequency with which cell-cell contacts (or proximity) were observed between two subclasses of cells. Two neighboring cells were considered contacting (or in proximity) if their centroid distance was <15 μm, which is approximately the size of the soma of a single cell in both human and mouse cortex (*31*). We determined how much this frequency was above random chance and how significant this difference was by comparing the observed contact frequency with the expected contact frequencies from spatial permutations. To avoid artifacts arising from the laminar organization of cells and spatial variation of cells density (namely, cell types in the regions of higher cell density or with similar laminar distributions can result in a higher contact frequency by random chance), we designed our spatial permutations to only disrupt the spatial relationship between neighboring cells while still preserving the laminar distribution and local density of each cell type (**Fig. S10**).

**Fig. 3.**
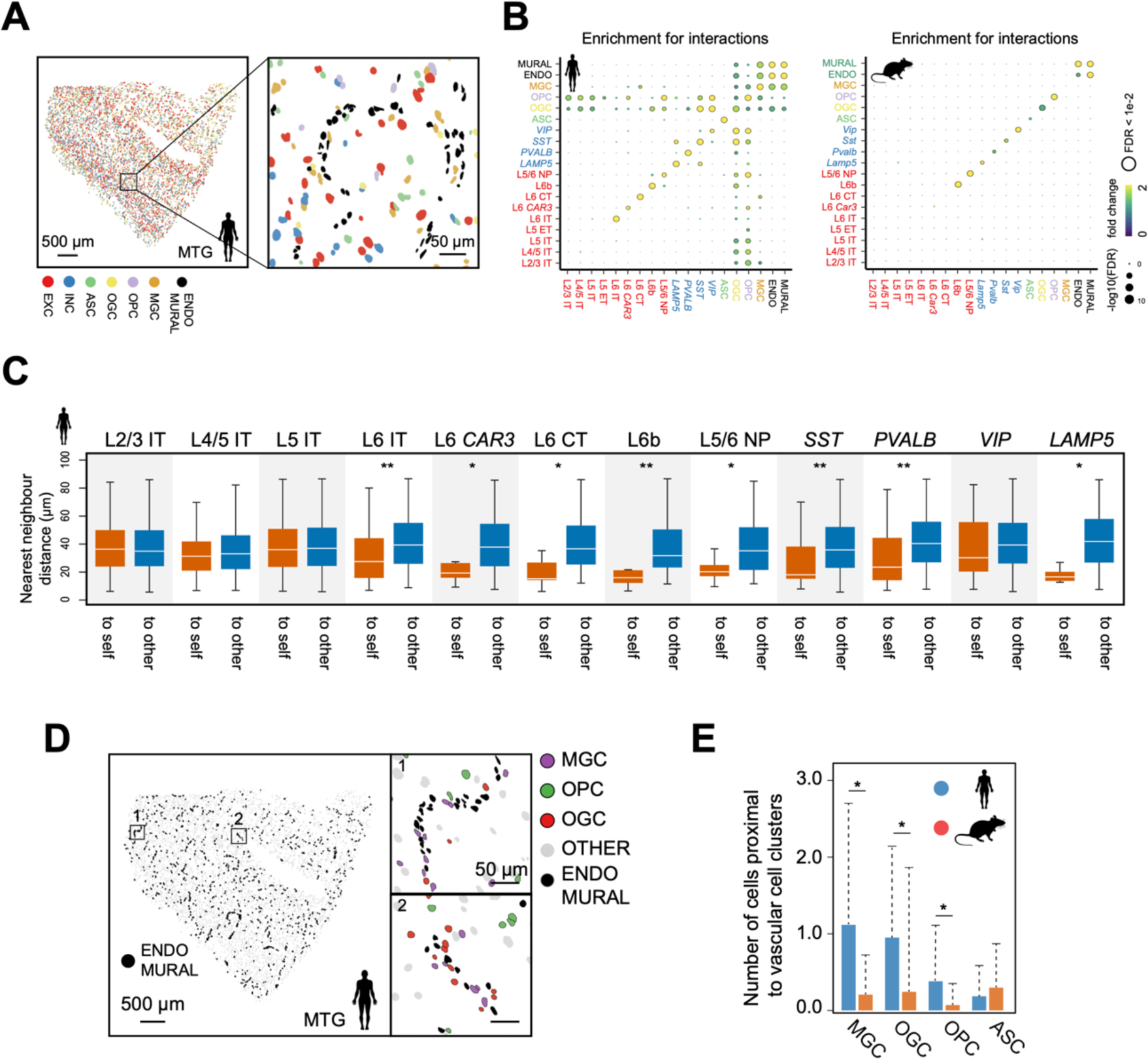
Cell-type-specific interactions in the human and mouse cortex. (**A**) Spatial map of excitatory and inhibitory neurons, as well as six major subclasses of non-neuronal cells in a human MTG slice (left) and a zoomed-in image of the boxed region (right). (**B**) Enrichment map of pairwise cell-cell interactions for subclasses of neuronal and non-neuronal cells in human (left) and mouse (right) cortex shown in dot plots. The enrichment was determined by comparing the observed interaction frequency between two types of cells with the distribution of the expected interaction frequencies generated by spatial permutations that disrupt the relationship between neighboring cells while preserving laminar distribution and local density of each cell type (**Fig. S10**). The color of the dot corresponds to the fold change between the observed interaction frequency and the average expected interaction frequency from the spatial permutations. The size of the dot corresponds to the significance level of the enrichment computed using one-tailed z-test. (**C**) Distribution of the nearest-neighbor distances between cells in individual neuronal subclasses to any cell from the same subclass (“to self”, orange) or to any cell from other subclasses (“to other”, blue). Boxplots shows the medians (middle white line), 25%-75% quantile (box), and 90% confidence interval (whiskers). The significance was determined by Wilcoxon rank-sum test (* P < 1e-3 and ** P < 1e-5). (**D**) Spatial map of vascular cells, including endothelial (ENDO) and mural (MURAL) cells for a human MTG slice (main panel). Zoomed-in images of boxed regions 1 and 2 with microglia, oligodendrocytes, and OPCs in colors, vascular cells in black, and other cells in grey are shown on the right of the main panel. (**E**) Average numbers of microglia, oligodendrocytes, OPCs, and astrocytes adjacent to each identified blood vessel in human (blue) and mouse (orange) cortices. Error bars are standard deviation. The significance was determined by Wilcoxon rank-sum one-tailed test (*P < 1e-3).

We observed interesting cell-type specific interaction patterns in the human cortex (**Fig. 3B** **and Fig. S11**). For neuron-neuron interactions, all four inhibitory neuronal subclasses (*LAMP5*, *PVALB*, *SST* and *VIP*) showed significant enrichment for self-interactions (**Fig. 3B, left**). Among the excitatory neurons, we observed enrichment for self-interactions within the subclasses in L6 (L6 IT, L6 IT *CAR3*, L6 CT, L6b, and L5/6 NP) (**Fig. 3B, left**). These results were further supported by examining the distributions of distances from individual neurons to their nearest neighbors in the same subclass or in different subclasses (**Fig. 3C**). Enrichment for self-interactions were also observed for the inhibitory neuronal subclasses in the mouse cortex (**Fig. 3B, right**), consistent with a previous finding (*32*), but the mouse cortex exhibited a lesser extent of self-interactions for the L6 excitatory neuronal subclasses (**Fig. 3B, right**).

We also found interesting human-specific enrichment for interactions among non-neuronal cells. For example, we observed enrichment for self-interactions among astrocytes in human (**Fig. 3B, left**), corroborated by independent observations made by electron microscopy (EM) reported in a recent bioRxiv preprint (*33*). It is surprising that the human cortex exhibited an enrichment for astrocyte self-interactions given that astrocytes are known to have non-overlapping territories in rodents (*34, 35*). Indeed, we did not observe substantial enrichment for astrocyte self-interactions in the mouse cortex (**Fig. 3B, right**). Likewise, we also observed enrichment for self-interactions among microglia in human but not mouse cortex (**Fig. 3B**).

Notably, we also observed substantially stronger enrichment for interactions of vascular cells (endothelial and mural cells) with oligodendrocytes and microglia in human as compared to mouse (**Fig. 3B**). Direct visualization of these cells in the human samples showed that many oligodendrocytes and microglia but not for astrocytes were clustered around and forming contacts with line-shaped vascular structures (presumably blood vessels) formed by endothelial and mural cells (**Fig. 3D**). These observations are also corroborated by the afore-mentioned EM study reported in bioRxiv (*33*) showing that oligodendrocyte and microglial cell bodies are adjacent to blood vessels whereas astrocytes contact blood vessels only with their process terminals but not cell bodies.

Interesting, such interactions between blood vessels and oligodendrocytes and microglia appeared to be human-specific and were rarely observed in the mouse samples (**Fig. 3B**). Quantifications of our MERFISH images showed that substantially more microglia, oligodendrocytes and OPCs, but not astrocytes, were contacting blood vessels in human than in mouse (**Fig. 3E** **and Fig. S12**).

The most remarkable cross-species differences on cell-cell interactions were observed between neurons and non-neuronal cells, in particular oligodendrocytes and microglia. We found significant enrichment for interactions between neurons and oligodendrocytes, including both mature oligodendrocytes and OPCs, in the human but not mouse cortex (**Fig. 3B**). Although somatic contacts between neurons and oligodendrocytes were also observed in the mouse cortex, the frequency of such events did not exceed the frequency expected from random chance. Moreover, a single neuron often formed contacts with several oligodendrocytes and/or OPCs in human, whereas such multi-way contacts were not enriched in mouse (**Fig. 4, A and B**). It is known that mature oligodendrocytes can be broadly classified into myelinating oligodendrocytes and non-myelinating perineuronal oligodendrocytes. The perineuronal oligodendrocytes in close proximity to the somata of neurons (*36*) can provide metabolic support to neurons (*37*). Our observed increase in interaction frequencies between perineuronal oligodendrocytes and neurons in the human cortex may thus be a result of evolutionary adaptation to higher energy demands during the firing of individual neurons in the human brain (*31*).

**Fig. 4.**
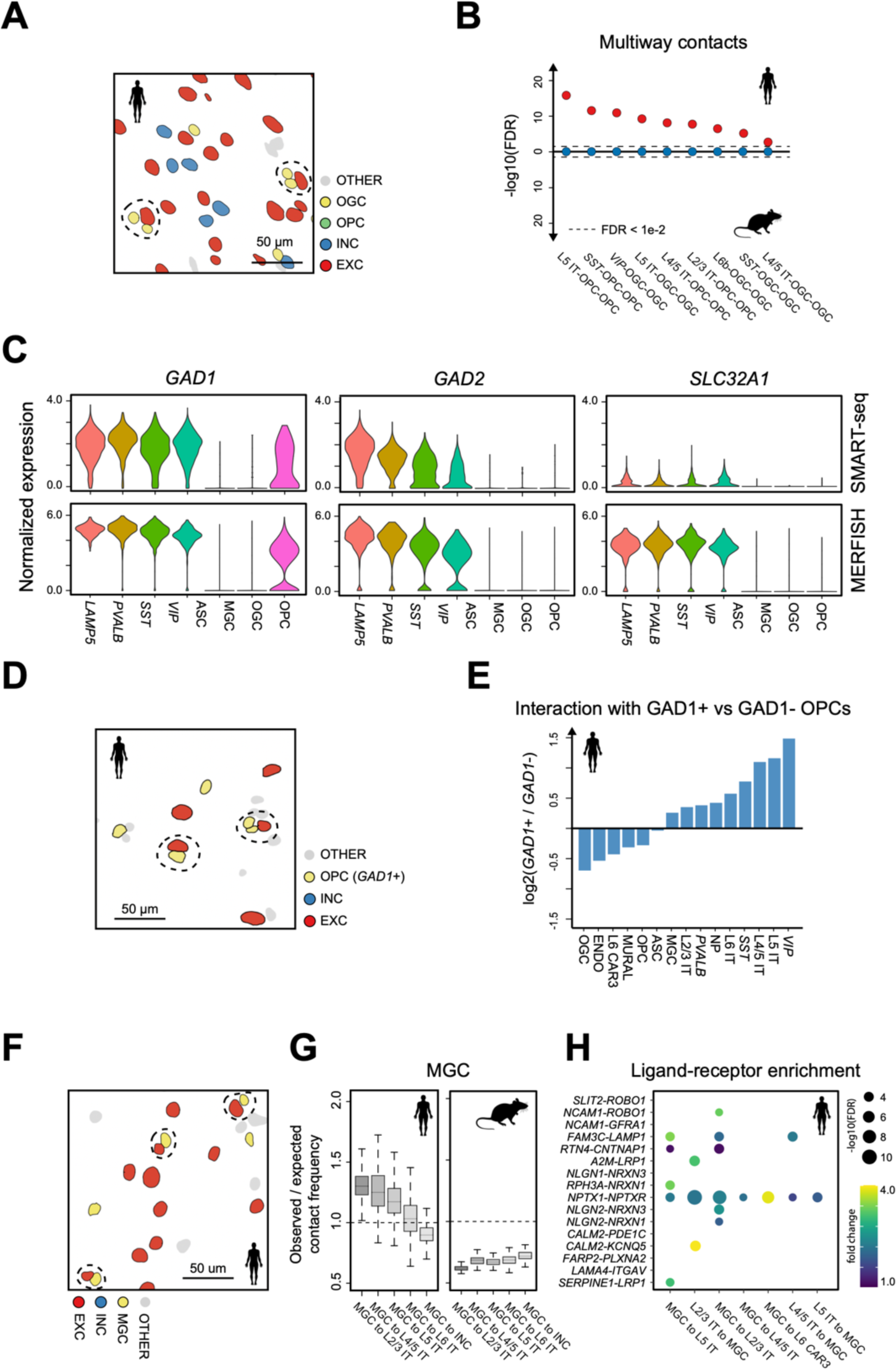
Cell-type-specific interactions between neurons and non-neuronal cells in the human and mouse cortex. (**A**) Spatial maps of excitatory neurons, inhibitory neurons, oligodendrocytes, and OPCs in an example region from a human MTG slice. Multiway contacts between neurons and oligodendrocytes are highlighted by dashed circles. (**B**) The significance level of multiway contacts between neurons and oligodendrocytes and/or OPCs in human (red circles) and mouse (blue circles) cortex. The significance level was determined by comparing the observed contact frequency with the expected frequencies from spatial permutations as described in **Fig. S10**. P-values were calculated using one-tailed z-test and then were adjusted to FDR (false discovery rate) using Benjamini and Hochberg procedure. (**C**) Normalized expression level of *GAD1, GAD2* and *SLC32A1* in seven subclasses of inhibitory neurons and glial cells in human MTG and STG determined by SMART-seq (top) and by 4,000-gene MERFISH (bottom). (**D**) Spatial maps of the excitatory neurons, inhibitory neurons, *GAD1*+ OPCs and *GAD1*-OPCs in an example region from a human MTG slice. Contacting OPCs and neurons are highlighted by dash line. (**E**) The log2 fold change of the contact frequency between *GAD1*+ OPC and other cell types over the contact frequency between *GAD1*-OPC and other cell types. (**F**) Spatial maps of excitatory neurons, inhibitory neurons, and microglia in an example region in a human MTG slice. Contacting excitatory neurons and microglia are highlighted by dash line. (**G**) Distribution of the ratio between observed and expected contact frequencies between microglia and L2/3 IT, L4/5 IT, L5 IT, L6 IT, and inhibitory neurons in human (left) and mouse (right) cortices. Expected contact frequencies were determined by the spatial permutations as described in **Fig. S10**. (**H**) Enrichment of ligand-receptor pairs in contacting microglia and IT neurons shown by dot plots. The size of the dot corresponds to the significance level and the color of the dot corresponds to the fold change of the observed ligand-receptor scores over their expected values (**Materials and Methods**).

Interestingly, Glutamate Decarboxylase 1 (*GAD1*), a gene encoding the enzyme that synthesizes GABA, was observed in a significant fraction of OPCs but not in mature oligodendrocytes (**Fig. 4C**). Our analyses of both MERFISH and SMART-seq data showed that ∼50% of the OPCs were *GAD1*+ in the human cortex, whereas Glutamate Decarboxylase 2 (*GAD2*) and the GABA transporter gene *VGAT* (*SLC32A1*) were not expressed in OPCs (**Fig. 4C**). Moreover, compared to *GAD1*-negative OPCs, *GAD1*-positive OPCs exhibited a substantially higher frequency to interact with neurons in human, including both inhibitory and excitatory neurons (**Fig. 4, D and E**).

Finally, our data revealed substantial human-mouse differences in interactions between neurons and microglia, the principal immune cells of the nervous system (*38*). Microglia play crucial roles in brain development and maintenance of proper neuronal function, and changes in microglial activity are linked to major human diseases (*7, 39*). Our MERFISH images showed that microglia frequently contacted neurons in the human cortex (**Fig. 4F**), likely representing satellite microglia (*40*). In addition, we found that these satellite microglia exhibited a higher frequency to interact with excitatory neurons than with inhibitory neurons in the human cortex (**Fig. 3B** **and** **Fig. 4, F and G**). Moreover, among excitatory neurons, microglia showed highest enrichment for their interactions with neurons in the upper layers, and the enrichment decreased with the cortical depth (**Fig. 4G**). In contrast, no significant enrichment in microglia-neuron interactions was observed in the mouse cortex (**Fig. 3B** **and** **Fig. 4G**).

To further explore the potential molecular mechanisms underlying the observed interactions between microglia and excitatory neurons, we identified ligand–receptor pairs enriched between contacting cells (**Materials and Methods**). Interestingly, several ligands and receptors that are genetically associated with neurodegenerative diseases were found to be enriched in contacting microglia and neurons (**Fig. 4H**). For instance, ligand-receptor pairs *CALM2-KCNQ5, NPTX1-NPTXR, A2M-LRP1, NCAM1-ROBO1, FAM3C-LAMP1, RTN4-CNTNAP1, NLGN2-NRXN3, NLGN2-NRXN1* were enriched in contacting L2/3 IT neurons and microglia (**Fig. 4H**), and among these Alpha-2-macroglobulin (*A2M*) is genetically associated with Alzheimer’s disease (*41*), low-density lipoprotein receptor-related protein 1 (*LRP1*) is a master regulator of tau uptake and spread (*42*), and neurexin 1/3 (NRXN1/3) are implicated in the pathophysiology of autism (*43*).

Because satellite microglia can maintain tissue homeostasis through phagocytosis of potentially toxic, extracellular protein aggregates in the healthy brain (*44, 45*), these microglia may function to protect nearby neurons. Indeed human genetics evidences suggest that microglia have a protective function that lowers the incidence of some neurodegenerative diseases (*46*). Our results thus suggest a potential functional interaction between microglia and excitatory neurons, especially those in the upper cortical layers, in the healthy human brain, and provide a potential mechanism that associates these ligand-receptors with neurodegenerative diseases: genetic mutations in ligand–receptors that disrupt the communications between microglia and neurons may leave nearby excitatory neurons unprotected and thus vulnerable in neurodegenerative disease. It also possible that accumulation of toxic aggregates in the diseased brain accrues these satellite microglia into inflammatory state that injures nearby excitatory neurons (*46*). These hypotheses are consistent with the observations that excitatory neurons in upper cortical layers are particularly vulnerable in neurodegenerative diseases (*47–49*).

In summary, using single-cell transcriptome imaging by MERFISH, we generated a molecularly defined and spatially resolved cell atlas of the human MTG and STG with high granularity for both neuronal and non-neuronal cells. The cell composition determined by MERFISH in these human cortical regions showed marked differences from that observed in several mouse cortical regions, with the human cortex exhibiting a five times higher glia-to-neuron ratio, three times higher inhibitory-to-excitatory neuron ratio, and five times higher IT-to-non-IT excitatory neuron ratio. Our MERFISH data showed both common and divergent features in the organizations of cells in human and mouse cortices. Most notably, we observed many differences in cell-cell interactions between human and mouse cortices, and the differences were particularly striking for interactions between neuronal and non-neuronal cells. The interactions between neurons and oligodendrocytes were significantly enhanced in the human cortex as compared to the mouse cortex. Moreover, microglial cells preferentially interacted with excitatory neurons over inhibitory neurons in the human cortex, whereas the mouse cortex did not exhibit enrichment for microglia-neuron interactions. Interestingly, ligand-receptor pairs genetically associated with neurodegenerative diseases were enriched in the interacting microglia-neuron pairs, suggesting possible molecular basis underlying the observed microglia-neuron interactions and connection of these cell-cell interactions to neurodegenerative diseases. It has been suggested that evolution of non-neuronal cells follows a more complex pattern than simply increasing the cell abundance, but additionally involves the diversification of glial cells (*50*). Our observations of the substantially enhanced interactions between neurons and non-neuronal cells in human as compared to mouse further expand upon this view. Given that such interactions are essential for normal functioning of the nervous system, we envision that MERFISH can be applied to study various diseases associated with non-neuronal cells to help uncover the molecular basis underlie their pathogenesis.

## Acknowledgments

We thank Jeff Ojemann (University of Washington) and Ryder Gwinn (Swedish Hospital in Seattle, WA) for providing human neurosurgical tissue samples and C. Dirk Keene and the staff in the University of Washington BioRepository and Integrated Neuropathology laboratory for their role in providing neurosurgical and postmortem tissue samples, and the Nancy and Buster Alvord Endowment to C. Dirk Keene. We thank Trygve Bakken, William Edward Allen and Cheen Euong Ang for helpful comments on the manuscript, and Rebecca Hodge and Julie Nyhus for assistance in postmortem sample preparation. This work is in part supported by the National Institutes of Health and Chan Zuckerberg Initiative, an advised fund of Silicon Valley Community Foundation. R.F. is a Howard Hughes Medical Institute Fellow of the Damon Runyon Cancer Research Foundation. X.Z. is a Howard Hughes Medical Institute Investigator.

## Supplementary Materials

### Materials and Methods

#### Ethical compliance

Surgical specimens were obtained from local hospitals (University of Washington Medical Center and Swedish Hospital in Seattle) in collaboration with local neurosurgeons. All patients provided informed consent and experimental procedures were approved by hospital institute review boards before commencing the study. De-identified postmortem human brain tissue was collected after obtaining permission from decedent next-of-kin. The Western Institutional Review Board (WIRB) reviewed the use of de-identified postmortem brain tissue for research purposes and determined that, in accordance with federal regulation 45 CFR 46 and associated guidance, the use of and generation of data from de-identified specimens from deceased individuals did not constitute human subjects research requiring institutional review board review.

Postmortem tissue collection was performed in accordance with the provisions of the United States Uniform Anatomical Gift Act of 2006 described in the California Health and Safety Code section 7150 (effective 1/1/2008) and other applicable state and federal laws and regulations. Tissue procurement from neurosurgical donors was performed outside of the supervision of the Allen Institute at local hospitals, and tissue was provided to the Allen Institute under the authority of the institutional review board of each participating hospital. A hospital-appointed case coordinator obtained informed consent from donors before surgery. Tissue specimens were de-identified before receipt by Allen Institute personnel. The specimens collected for this study were apparently non-pathological tissues removed during the normal course of surgery to access underlying pathological tissues. Tissue specimens collected were determined to be non-essential for diagnostic purposes by medical staff and would have otherwise been discarded.

#### Human brain processing

Human brains were transported on ice and processed as bisected hemisphered by embedding in Cavex Impressional alginate (Cavex Life Cast Normal Aglinate) prepared according to manufacturer’s instructions. Each cerebral hemisphere was then cut into 1 cm slabs and individually frozen in an isopentane and dry ice slurry. Each slab is then vacuum sealed and stored at -80 °C. MTG and STG samples were taken by allowing selected slabs to equilibrate to -20 °C for 2 hours. The region of interest was identified and dissected from the slab at -20 °C using a scalpel, vacuum sealed and stored at – 80 °C.

#### Animals

Adult C57BL/6 male mice aged 57-63 days were used in this study. Animals were maintained on a 12 hour: 12-hour light/dark cycle (2pm-2am dark period) with ad libitum access to food and water. Animal care and experiments were carried out in accordance with NIH guidelines and were approved by the Harvard University Institutional Animal Care and Use Committee (IACUC). Mice were euthanized with CO2, and their brain were quickly harvested and cut into hemispheres and each hemisphere was frozen immediately on dry ice in optimal cutting temperature compound (Tissue-Tek O.C.T.; VWR, 25608-930), and stored at -80 °C until cutting. Frozen brain hemispheres were sectioned at -20 °C on a cryostat (Leica CM3050 S). Slices were removed and discarded until the visual cortex region (VIS) was reached. Two 10-µm-thick slices were cut from anterior to posterior from each mouse and were placed onto salinized coverslips for imaging. In total two slices were collected for each animal and four slices were imaged.

#### MERFISH gene selection

To identify transcriptionally distinct cell populations in the human MTG and STG with MERFISH, we designed a panel of 4000 genes. In details, we first use a combination of two approaches, mutual information (MI) analysis and differentially expressed (DE) gene analysis, to select marker genes based on previously published single nucleus SMART-seq data in the MTG of the human brain (*8*) as described previously (*15, 16*). In the first approach, we selected the top 300 genes with the greatest MI values for excitatory neuronal clusters and inhibitory neuronal clusters, respectively. In the second approach, we included all DE genes for each cluster when compared to the remaining cells in its respective group, namely excitatory or inhibitory neuron group. This yielded 486 DE genes for excitatory neuronal clusters and 599 DE genes for inhibitory neuronal clusters. We then removed overlapping genes among these 4 groups of MI and DE genes, resulting in a total of 1,233 marker gene candidates. Next, we then added the manually and computationally selected genes provided by the Chan Zuckerberg Initiative SpaceTx Consortium and removed the genes that overlap with the 1,233 genes deriving from MI analysis and DE analysis, resulting in 1,680 maker gene candidates. To finalize the maker gene list, we further remove genes that were either not long enough to construct 60 target regions (each 30-nucleotide (nt) long) without overlap or whose express levels were outside the range from 0.01 to 500 FPKM, as measured by bulk RNA-seq. This yielded 764 marker genes. For the remaining 3,236 genes, we randomly selected genes based on three criteria: 1) the genes were long enough to construct 60 target regions without overlap; 2) the express levels of the genes were within the range of 0.01 to 500 FPKM; 3) the genes did not overlap with 764 cell type marker genes.

#### Design and construction of the MERFISH encoding probes

Each RNA species was encoded with a unique binary word from a set of 48-bit, Hamming Distance 4 (HD 4), Hamming Weight 4 (HW 4), binary barcodes (**Table S1**). This codebook contains 4,140 unique barcodes and every barcode was separated by a Hamming distance of at least 4 from all other barcodes. Each of 4,000 selected RNA species was randomly assigned a unique binary barcode from all 4,140 possible 48-bit barcodes. The remaining unassigned 140 barcodes were left as blank controls for misidentification quantification.

For each RNA species, we designed 60 non-overlapping encoding probes to imprint the selected binary codeword onto the corresponding transcript, each encoding probe containing a 30-nt target sequence complementary to one of the 30-nt target regions.

The encoding regions were identified using an algorithm as described previously (*51*). In brief, we selected target regions that had a GC fraction between 30% and 70%, a melting temperature TM within the range of 60-80 °C, an isoform-specificity index between 0.75 and 1, a gene-specificity index between 0.75 and 1, and no homology longer than 15 nt to rRNAs or tRNAs. Each of the 48 bits was then assigned a 20-nt three-letter readout sequence. Afterwards, each encoding probe was constructed to contain a unique 30-nt target region and three of the four 20-nt readout sequences corresponding to the four ‘1’ bits assigned to the transcript. The encoding probes additionally contained two priming regions for amplification: a 20-nt primer binding region at the 5’ end and the reverse complement of the T7 promoter at the 3’ end as previously described (*51*). The encoding probe libraries were purchased from Twist Biosciences and amplified as previously described (*51*). The encoding probe sequences for 4,000 genes are listed in **Table S2**. To test whether the molecular crowding associated with imaging 4,000 genes caused a substantial reduction in the detection efficiency, we also performed MERFISH imaging on 250 genes chosen from the 764 marker genes. The codebook for 250-gene is listed in **Table S3** and encoding probe sequences for 250 genes are listed in **Table S4**.

#### Design and construction of the MERFISH readout probes

In this study, 48 readout probes complementary to the 48 readout sequences were designed, each corresponding to one of the 48 bits in the barcodes. Readout probes were conjugated to one of the dye molecules (Alexa750, Cy5 and Atto565/Cy3B) via a disulfide linkage, as described before (*51*). These readout probes were synthesized and purified by Biosynthesis, Inc., resuspended immediately in Tris-EDTA (TE) buffer, pH 8 (Thermo Fisher) to a concentration of 100 μM and stored at -20 °C. The readout sequences used in this study are listed in **Table S5**.

#### Sample staining

Tissue slices were allowed to briefly thaw on coverslips at room temperature, and then were fixed by treating with 4% PFA in 1×PBS for 10-15 minutes. We then washed coverslips three times with 1×PBS and stored them in 70% ethanol at 4°C for overnight to permeabilize cell membranes. After permeabilization, tissue slices were exposed under the multi-band light emitting diode arrays for three hours to eliminate the autofluorescence background (*18*). To perform MERFISH encoding probe staining, samples were incubated for 30 min in encoding wash buffer comprising 2× SSC (Ambion), 30% (vol/vol) formamide (Ambion), and 0.1% (vol/vol) murine RNase Inhibitor (New England Biolabs). A drop of 60 μL of 4,000 gene encoding probes (∼1 nM per encoding probe, 60 probes per gene) and acrydite-modified poly(dT) locked nucleic acid (LNA) probes with a unique 20-nt readout sequence (2 μM) in encoding hybridization buffer was added to a parafilm in a 150mm-diameter dish and were covered with cell-containing coverslips. Encoding hybridization buffer was comprised of encoding wash buffer supplemented with 0.1% (wt/vol) yeast tRNA (Thermo Fisher), 0.1% (vol/vol) murine RNase Inhibitor (New England Biolabs), and 10% (wt/vol) dextran sulfate (Sigma). The samples were then incubated in a cell culture incubator at 37 °C for ∼48h, followed by washing with encoding wash buffer twice at 47 °C for 30 min. The samples then were incubated for 10 min with a 1:500 dilution of 0.12-μm-diameter light yellow beads (Spherotech, FP-0245-2), which were used as fiducial markers to align images obtained from sequential rounds of hybridization.

#### Sample expansion

To expand the samples labeled with MERFISH encoding probes, we adopted a previously published gel embedding and expansion protocol (*14, 20, 52*). Briefly, the samples were first incubated for 10 min in degassed monomer solution consisting of 2 M NaCl, 7.7% (wt/vol) sodium acrylate (Sigma), 4% (vol/vol) of 19:1 acrylamide/bisacrylamide, 60 mM Tris-HCl pH 8 and 0.2% (vol/vol) TEMED (for buffer exchange). The remaining monomer solution was then kept on ice and further added 0.2% (wt/vol) ammonium persulfate just before casting the expansion gel.

To cast the gel, 25 µL of the ice-cold monomer solution containing 0.2% (wt/vol) ammonium persulfate was added to the surface of a glass plate that had been treated with GelSlick (Lonza). Coverslips containing the encoding-probe-labeled slices were dried quickly with KimWipes (Kimtech) and inverted onto the 25 µL droplet to form a uniform thin layer of gel solution between the coverslip and the glass plate. The sandwiched coverslip and glass plate with gel solution were then transferred to a nitrogen-filled chamber for 2 hours to complete gel polymerization. The coverslip and the glass plate were then separated by using a thin razor blade. The coverslips with expansion gels were transferred to a digestion buffer, which was made up of 2% (wt/vol) Sodium dodecyl sulfate (SDS) (Thermo Fisher), 0.5% (vol/vol) Triton X-100 and 1% (vol/vol) Proteinase K (New England Biolabs) in 2× SSC. The samples were digested in digestion buffer overnight in a 37 °C incubator. After digestion, samples were expanded in 1 × SSC buffer supplemented with 0.2% (vol/vol) Proteinase K at room temperature. During expansion, the buffer was changed every 20 minutes for three times.

To stabilize the expanded gel for sequential rounds of readout probe hybridization and imaging, we re-embedded the expansion gel in a nonexpandable polyacrylamide gel. Briefly, expanded polyacrylite gel was buffer exchanged in re-embedding solution comprised of 4% (vol/vol) of 19:1 acrylamide/bis-acrylamide, 100 mM NaCl, 60 mM Tris⋅HCl pH 8 and 0.2% (vol/vol) of TEMED for 20 min. The remaining re-embedding solution was kept on ice and further supplemented with 0.2% (wt/vol) ammonium persulfate just before casting the nonexpandable polyacrylamide gel. The expanded polyacrylate gels were transferred to silanized coverslips prepared as previously described, dried with KimWipes and added 100 μL re-embedding gel solution with ammonium persulfate. The samples were then put in a nitrogen chamber, covered with a GelSlick treated glass plate to facilitate polyacrylamide gel polymerization at room temperature for 1 h. The coverslip and the glass plate were again separated by using a razor. The samples were either used for MERFISH measurement immediately or stored in 2 × SSC containing 0.1% (vol/vol) murine RNase inhibitor at 4°C for no longer than a week.

#### MERFISH imaging

MERFISH imaging were carried out on the imaging platform as described previously (*14*). In the first round we imaged the cell nucleus stained with DAPI and poly-dT labeled readout probe complementary to the poly-dT LNA anchor probe. We then performed 16 rounds of three-color MERFISH imaging for the both 4,000- and 250-gene measurements. Both DAPI and poly-dT were imaged at six focal planes separated by 1 μm in z. Each MERFISH round consisted of readout probe hybridization (10 min), washing (5 min), and imaging of ∼1,000 tiled fields of view (FOV) (220 μm x 220 μm per FOV), readout fluorophore cleavage by TCEP (15 min), and rinsing with 2 x SSC (5 min). For each round, images were acquired with 750-nm, 650-nm, and 560-nm illumination at six focal planes separated by 1 μm in z to image the readout probes in addition to a single z-plane image with 405-nm illumination to image the fiducial beads on the glass surface for image registration. Specifically, the readout probes staining was done by flowing 10 nM (each) readout probes in hybridization buffer composed of 2 × SSC, 10% (vol/vol) ethylene carbonate (Sigma-Aldrich, E26258), 0.1% (vol/vol) murine RNase inhibitor (NEB), 0.5% (vol/vol) Triton X-100 and 0.4% (vol/vol) TWEEN 20 in nuclease-free water, following by washing with a wash buffer containing 2× SSC and 10% (vol/vol) ethylene carbonate in nuclease-free water. TWEEN 20 can passivate coverslip surface and reduce non-specific readout probe binding. Readout probe imaging was then performed in imaging buffer containing 5 mM 3,4-dihydroxybenzoic acid (Sigma), 2 mM trolox (Sigma), 50 µM trolox quinone, 1:500 recombinant protocatechuate 3,4-dioxygenase (rPCO; OYC Americas), 1:500 Murine RNase inhibitor, and 5 mM NaOH (to adjust pH to 7.0) in 2×SSC. After imaging, dyes on the readout probe were removed by flowing in cleavage buffer comprising 2 × SSC and 50 mM of Tris (2-carboxyethyl) phosphine (TCEP; Sigma) to cleave the disulfide bond connecting dyes to the probes. The imaging buffer and hybridization buffer were changed every 12 hours to prevent the decay of imaging quality.

#### MERFISH decoding

To align the 3-dimensional x, y, z image stacks for all 48-bit MERFISH measurements, we calculated the offset between the corresponding high-pass filtered fiducial bead images with subpixel resolution by finding the peak of the image cross-correlation and applied these offsets to align the image stacks. To identify the RNA transcripts in the registered image stacks, we modified our previously published pixel-based decoding algorithm (*14, 51*). Our decoding framework included two stages: crude decoding and fine decoding. Each field of view (FOV) contained six z-planes and decoding between z-planes were independent. In the crude decoding, we first randomly selected one z-plane from each of the 100 randomly selected FOVs for pixel-based decoding. In short, we assigned barcodes to each pixel independently, then aggregated adjacent pixels with the same barcodes into putative RNA molecules and filtered the list of putative RNA molecules to enrich for correctly identified transcripts (*14*). In detail, to assign each pixel to one of the 4,140 (4,000 coding and 140 blank) barcodes, we compared the 48-dimensional intensity vectors measured for each pixel to the vectors corresponding to the 48-bit valid barcodes. To aid comparison, we first normalized intensity of each image by its median intensity across the 100 random FOVs (using the randomly selected z-plane for each FOV) to eliminate the intensity variation between hybridizations and color channels. After intensity normalization, we further normalized intensity variations across pixels by dividing the 48-dimensional intensity vector for each pixel by its L2 norm, such that the magnitude in the 48-dimensional space was equal to 1 for each pixel. We similarly normalized each of the 4,140 valid 48-bit barcodes, such that the magnitude in the 48-dimensional space was equal to 1 for each valid barcode. To assign a barcode to each pixel, we identified the normalized barcode vector that was closest to the pixel’s normalized intensity vector in the 48-dimensional space. We excluded pixels with distance larger than 0.65 away from any valid barcode in the first step. To further reduce the chance of transcript misidentification, for each remaining pixel, we calculated two properties: the mean intensity, referred to as “pixel intensity”, and the distance between the normalized intensity vector of the corresponding pixel to the closest valid barcode, referred to as “pixel distance” hereafter. We anticipated that incorrectly identified barcodes would likely have a dimmer mean pixel intensity and a larger distance as compared to correctly identified barcodes. To distinguish the valid pixel from the misassigned ones, we next trained a support vector machine (SVM) using pixels assigned to blank barcodes as negative control and pixels assigned to RNA-coding barcodes as positive ones. Because the RNA-coding pixels usually far outnumber the blank pixels, we down sampled the RNA-coding pixels to the same number as the blank pixels to avoid sample imbalance. Using the trained SVM, we next predicted the probability of a pixel being a valid (positive) barcode. As expected, the pixels with dimmer intensity or large distance were predicted with lower probability while bright pixels with small distance were assigned with a larger probability.

In the stage of fine decoding, we used the SVM classifier trained, and scaling factors estimated during the crude decoding to decode the rest of the z-planes. We first filtered any pixels that have a distance greater than the distance cutoff (0.65). For the remaining pixels in each z-plane, we then assigned barcode ID to each pixel and grouped adjacent pixels if they were assigned to the same barcode ID to create a list of putative RNA molecules. We measured the confidence of the identified molecules by calculating the probability of at least one pixel is valid as following:

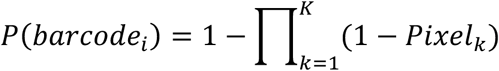

In which *K* is the total number of grouped adjacent pixels assigned to *barcode_i_* for this molecule and *Pixel_i_* represents the probability of pixel *k* is assigned to *i*-th barcode *barcode_i_* determined by SVM classifier. To avoid overfloating, instead of calculating the probability, we computed the log likelihood of the barcode:

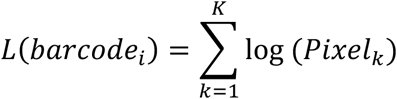

We next ranked all identified barcodes based on the barcode likelihood *L*(*barcode_i_*) and determined the likelihood cutoff that resulted in a gross barcode misidentification rate of 5%, where the gross barcode misidentification rate was estimated as (the mean count per blank control barcode) / (the mean count per coding barcode).

#### MERFISH image segmentation

We performed cell segmentation using co-staining of DAPI and total polyA RNA as demonstrated previously for MERFISH analysis (*15*). Using a deep learning-based cell segmentation algorithm (cellpose) (*21*), we first segmented nuclei with the DAPI image with diameter parameter of 150 pixels in the “nuclei” mode. We segmented individual z-planes for each FOV. We then identified the centroid position of each segmented nucleus in each z-plane and connected the centroids across different z-planes if they were within distance of 30 pixels (∼3.2 μm) in xy direction, representing the same nucleus imaged in different z-sections. Because each of these image operations was applied to the full z-stack of images for each FOV, the generated nuclei segmentation boundaries were naturally in 3D. This algorithm was able to largely address the situation in which two adjacent cells partially overlapped along the xy axial direction.

We next segmented soma by total polyA RNA image also using cellpose with diameter parameter of 200 in a “cytoplasm” mode. We then used the segmented nuclei as seeds and segmented cell bodies as a mask on image manifolds established by total polyA RNA image and performed watershed algorithm. In this manner, we assigned unique IDs for each cell body based its unique the nuclear seed. Nuclei and cell bodies were assigned consistent cell IDs for downstream analysis. We performed watershed for individual z planes for each FOV to create the 3D segmentations.

#### Cloud-based massively parallel MERFISH decoding

Because MERFISH decoding is independent between z-planes, therefore, the decoding procedure can be divided into thousands of small independent tasks. This allowed us to speed up decoding of large-scale MERFISH datasets by leveraging the power of cloud computing platform. We set up an auto-scale SLURM cluster on Google cloud platform by following the instruction (https://cloud.google.com/architecture/deploying-slurm-cluster-compute-engine). The SLURM cluster had access to two types of compute engines including n1-standard-4 (4 vCPUs, 15 GB memory) and n1-standard-16 (16 vCPUs, 64 GB memory), up to 1,000 vCPUs and 2TB Standard persistent disk. n1-standard-4 machines with limited memory was used for decoding thousands of z-planes in parallel. n1-standard-16 was used for more memory-consuming tasks such as creating global mosaic image and barcode filtering. In both cases, preemptible nodes were used to reduce the cost. The decoding pipeline was managed using Snakemake - a tool to create reproducible and scalable data analyses (*53*). MERFISH data was stored in the Google bucket (STANDARDED) with unlimited storage that allowed the SLURM cluster to access.

#### Unsupervised clustering of 4000-gene MERFISH data

Because the cell type marker genes were identified using single nucleus SMART-seq data (*8*), in this study, we used the RNA molecules identified in the nucleus for clustering analysis. With the nucleus-by-gene matrix, we first preprocessed the matrix by several steps. The segmentation approach described above generated a small fraction of putative “nuclei” with very small total volumes or low RNA abundance due to segmentation artifacts, as well as some cells that overlapped in the 3-D dimension and were not properly separated. We first filtered any cells with segmentation appeared in less than three z-planes to avoid spurious segmentations. Second, we filtered cells whose centroid was within 100 pixels to the edge of the FOV to avoid edge effect. Third, we removed the segmented “cells” that had a volume that was either less than 300 µ*m*^2^ or larger than 6000 µ*m*^2^. Fourth, we calculated the RNA molecule density in each nucleus and filtered instances with density less than 0.1 molecule / µ*m*^2^ which was closed to the molecule density outside the cell body, likely representing “empty” nucleus introduced by spurious segmentations or out-of-focus signal.

Using the remaining nucleus, we next performed clustering analysis. In detail, we first normalized the nucleus-by-gene count matrix using “scTransform”, a modeling framework for the normalization and variance stabilization of molecular count data from scRNA-seq experiments (*54*). To remove the differences in RNA counts due to the incompleteness of nucleus, we further normalized the RNA counts per cell by regressing out the imaged volume of each cell using “scTransform” by setting the “vars.to.regress” to the area size of the nucleus. Next, we performed dimensionality reduction using principal component analysis (PCA), restricted to the 30 principal components (PCs) with the highest eigenvalues, and finally visualized using a 2D UMAP embedding (*55*). In the UMAP space, the batch effect between MERFISH experiments was further eliminated using Canonical Correlation Analysis (CCA) which was implemented in Seurat V3 (*9*). To identify transcriptionally distinct cell clusters, we performed graph-based Leiden (*56*) community detection (k=15; resolution=0.5) in the 30 PCs-space.

We first annotated identified clusters to major cell types based on the expression of canonical marker genes. Next, we determined the excitatory and inhibitory neuronal clusters by joint analysis of snSMART-seq and MERFISH. In detail, we performed normalization using “scTransform” for MERFISH and SMART-seq independently. ScTransform automatically selected 2,000 variable genes in both datasets as “anchors” for integration. Second, we used the selected anchors and CCA to generate the joint low-dimensional embedding space between MERFISH and SMART-seq. In the joint space, we identified clusters using graph-based clustering method (Leiden). A key parameter in Leiden is resolution, a higher resolution usually resulting in more clusters. How to choose the optimal resolution is described with more details below. First, for a clustering result generated by a given resolution, we trained a KNN classifier in the joint embedding space on 80% of the dataset and then estimated the prediction accuracy on the remaining 20%. We conducted this five times (also known as five-fold cross validation) to estimate the averaged prediction accuracy. As expected, we observed a decrease in prediction accuracy with increasing resolution, suggesting “over splitting” of clusters. In this study, we chose the resolution that yield 3% misidentification rate. To avoid the errors introduced by the integration algorithm, we next filtered any “outlier” clusters that were mostly identified by one dataset. In detail, for each cluster, we calculated the ratio between cell proportion determined by SMART-seq and MERFISH and then normalized these ratios to z-score across all clusters. We removed any clusters whose z-score was larger than 3 or less than -3. Finally, for each cluster, we performed differentially expression (DE) analysis using Wilcoxon Rank Sum test in MERFISH and SMART-seq data separately (Pvalue < 1e-2; fold-change > 1.2). A cluster which failed to find a DE gene in both MERFISH and SMART-seq was merged with the closest clusters in the joint embedding space. This was repeated until no clusters could be merged anymore.

#### Supervised classification of 250-gene MERFISH data

To make use of our 250-gene MERFISH datasets, we performed supervised classification to predict the cluster labels for cells measured in the 250-gene experiments based on the annotation of the 4,000-gene MERFISH datasets. We performed normalization using “scTransform” for 4,000-gene and 250-gene MERFISH dataset separately. In this case, ScTransform selected 2,000 variable genes for 4,000-gene MERFISH dataset and 250 variable genes for 250-gene MERFISH datasets.

Second, we used CCA to generate the joint low-dimensional embedding space between 4,000- and 250-gene datasets. We next applied the neighborhood-based classifier to make a classification of labels for cells in 250-gene dataset. Briefly, we assigned a cluster label to each cell in 250-gene dataset based on the cluster labels of its neighboring cells in the annotated 4,000-gene dataset. The contribution of the neighboring cells in the 4,000-gene dataset was weighted by their distances to the query cell in the 250-gene dataset. In detail, for each cell ***c*** in 250-gene dataset, we identified the nearest ***s*** (|***s***|=50) cells in the 4,000-gene dataset in the joint embedding space in the first 30 dimensions. For each cell *s_i_* in ***s***, we calculated the Euclidean distance dist(*c*, *s_i_*) between the query cell ***c*** and reference cell *s_i_*. This distance was then weighted based on the distance to the *s*-th reference cell in the 4,000-gene dataset:

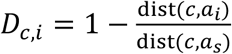

The weighted distance was converted to weighted similarity:

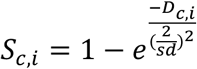

where *sd* is set to 1 by default. Finally, we normalized the similarity across all ***s*** anchors:

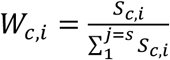

Hence *W_c,i_* is the weighted similarity between query cell ***c*** and reference cell *s_i_*.

Using the weighted similarity, we next predicted the cluster label for query cell ***c***. In detail, let **L** ∈ ℛ*^k×3^* be the binary label matrix for *k* cells with *t* clusters. *L_i,j_* = 1 indicates the class label for *i*-th 4,000-gene cell is *j*-th cluster. The row sum of **L** must be 1, suggesting each 4,000-gene cell can only be assigned to one cluster label. We then compute label predictions for query cells as **P***^l^*:

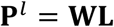

The resulting **P**^5^ is a probability matrix, 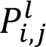 indicates the probability of a cell *i* assigned to cluster *j*. Each cell in the 250-gene measurement was assigned to the cluster label that had the maximum probability. Cells with the maximum assignment probability less than 0.4 was removed from downstream analysis.

#### Cell-cell interaction analysis

We defined two cells to be contacting or interacting if they were within 15 μm of each other, a distance that was close to the size of a soma for both human and mouse. We calculated the interaction frequency between two cell types (i.e. A and B) by counting the number of cell pairs between A and B whose distance were below this 15 μm threshold in the 2D spatial map. We noticed and corrected the following biases in the interaction map. First, the cells in the regions of high cell density usually resulted in higher interaction frequency as compared to those in the sparse regions. Second, cell types of similar laminar organization resulted in higher self-interaction frequency. To correct for these biases, we randomized the cell position within a short-range distance to disrupt the proximity between neighboring cells while largely preserves the higher-order laminar organization and local cell density (**Fig. S10**). In detail, we randomized the position of each cell within a radius of 50 μm of its measured position and determined the interaction frequency between any two cell types, and we performed such spatial permutation 1,000 times to generate a null distribution. By calculating the z-score for the observed interaction frequencies based on the null distribution, we obtained a P value for the significance of interaction between two cell types. Finally, P-values were corrected using Benjamini & Hochberg (BH). To examine the effect of permutation radius, we performed the spatial permutation test with different radii from 35 μm to 100 μm and observed similar results (**Fig. S11**).

#### Ligand-receptor analysis

To determine the enrichment of a ligand–receptor pair (*L_i_*-*R_i_*) in two interacting cell types (i.e. A and B), we first identified cells pairs between A and B that were contacting as described in the section above. For each interacting cell pairs, we calculated the product of the normalized expression (logCPM) of the ligand (*L_i_*) by one cell and receptor (*R_i_*) by another cell, and determined the sum of this product across all interacting cell pairs as the ligand-receptor score for A and B. We then determined the enrichment significance using empirical shuffling. In each shuffling, we randomly selected the same number of cell pairs from A and B whose distances were larger than 30 μm and calculated the ligand-receptor score for *L_i_* − *R_i_* as described above. We repeated this for 1,000 times to obtain the null distribution. Using one-tailed z-test, we obtained a P value for the significance. Finally, P values were adjusted using Benjamini & Hochberg (BH).

---

**Fig. S1.**
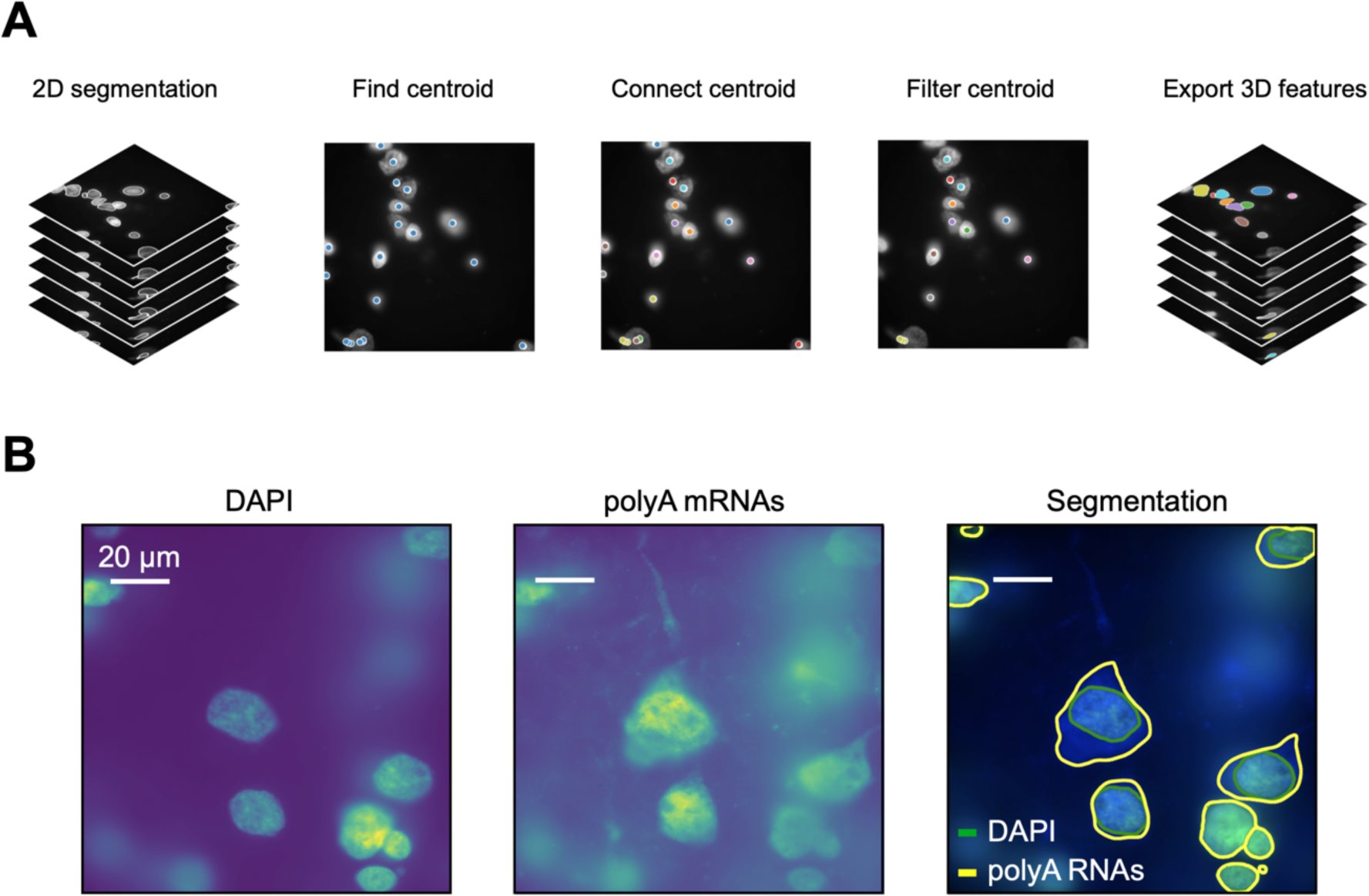
Cell segmentation based on DAPI and total polyA RNAs signals. (**A**) The schematic of 3D cell segmentation. Cell boundaries first identified in each z-plane separately using deep learning-based segmentation algorithm (Cellpose) (*21*). The centroids of each 2D segmentations were connected between z-planes if their distance in xy was smaller than 2 μm. The connected centroids represent the same cell sampled in different z-planes. Cells that did not appear in at least two consecutive z-planes were likely due to segmentation errors and filtered out. (**B**) Segmentation of cells with DAPI and total polyA RNA co-stains in one field-of-view. DAPI and total polyA RNA images of a single field-of-view are shown in the left and middle panels, respectively, and the right panel shows the segmentation boundaries for each cell and each nucleus marked in yellow and green, respectively.

**Fig. S2.**
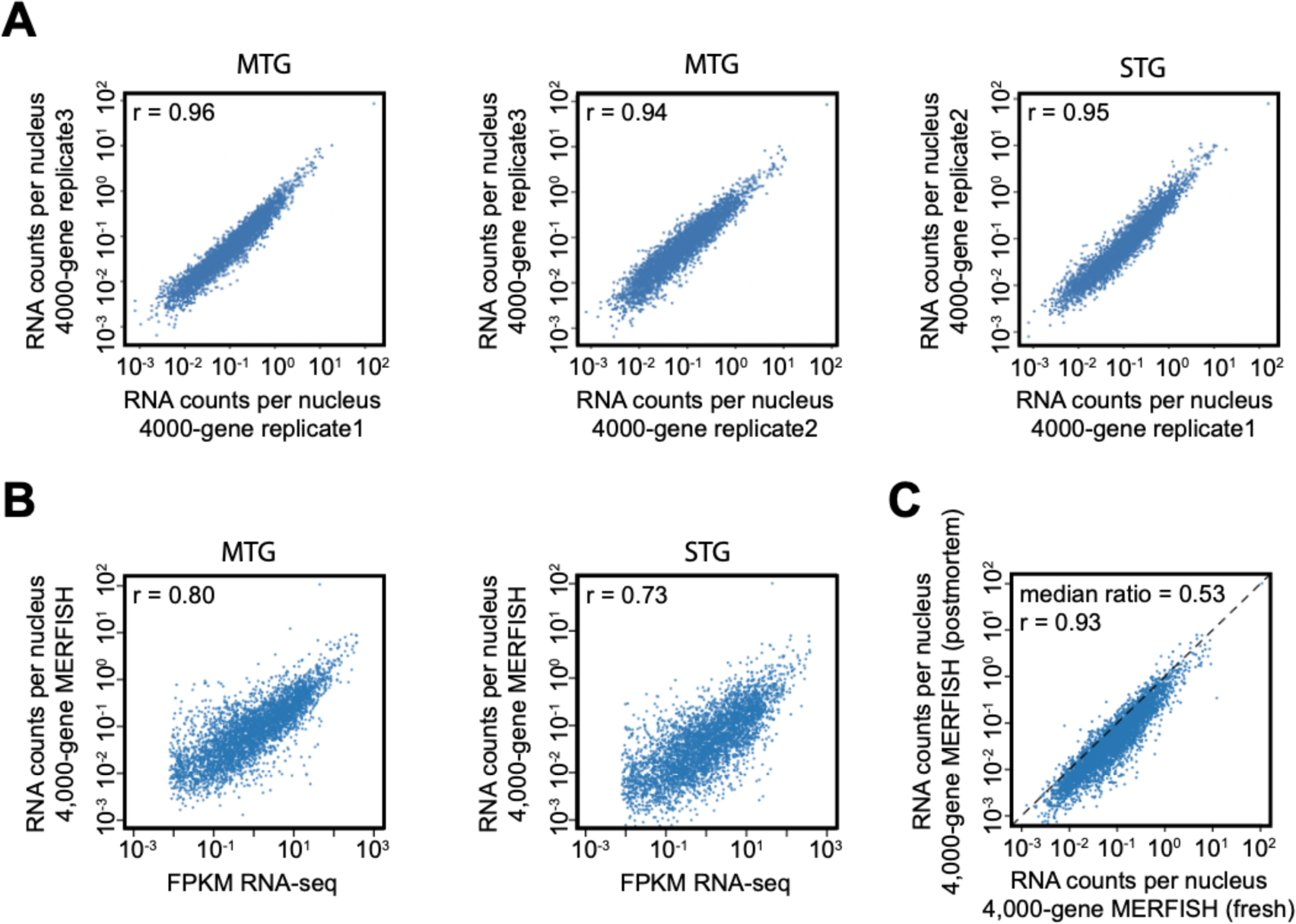
Reproducibility of MERFISH data between biological replicates and correlation of MERFISH data with bulk RNA sequencing results. (**A**) The scattered plots of the RNA copy number of individual genes per nucleus detected in the 4,000-gene MERFISH measurements between biological replicates in the human MTG and STG slices. (**B**) The RNA copy number of individual genes per nucleus detected in the 4,000-gene MERFISH measurements of MTG (left) and STG (right) versus the FPKM from bulk RNA-seq. (**C**) The average RNA copy number of individual genes per nucleus detected in the 4,000-gene MERFISH measurements of the fresh-frozen, neurosurgical MTG samples versus that of the postmortem STG samples. Three MTG biological replicates and two STG biological replicates were respectively used to determine the average RNA copy numbers. r stands for the Pearson correlation coefficient.

**Fig. S3.**
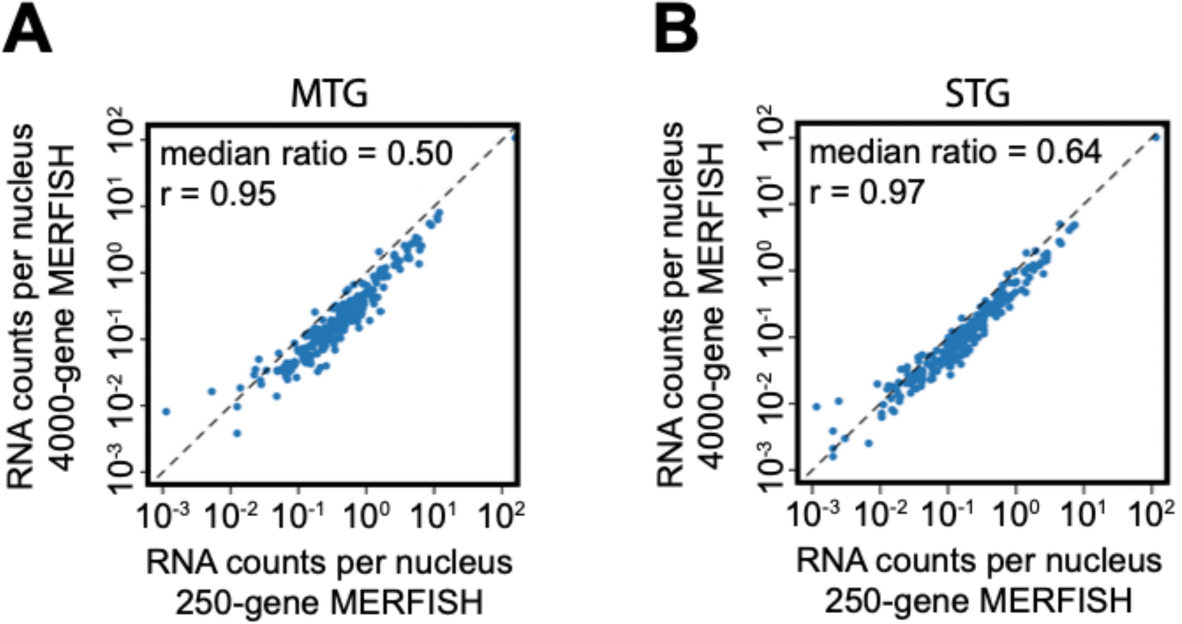
Comparison of 4000-gene and 250-gene MERFISH data. The average RNA copy number of individual genes per nucleus detected in 4,000-gene measurements are compared to that in the 250-gene measurements for the 250 genes that are shared in both measurements. (**A**) The comparison for the MTG samples. (**B**) The comparison for the STG samples. r stands for the Pearson correlation coefficient.

**Fig. S4.**
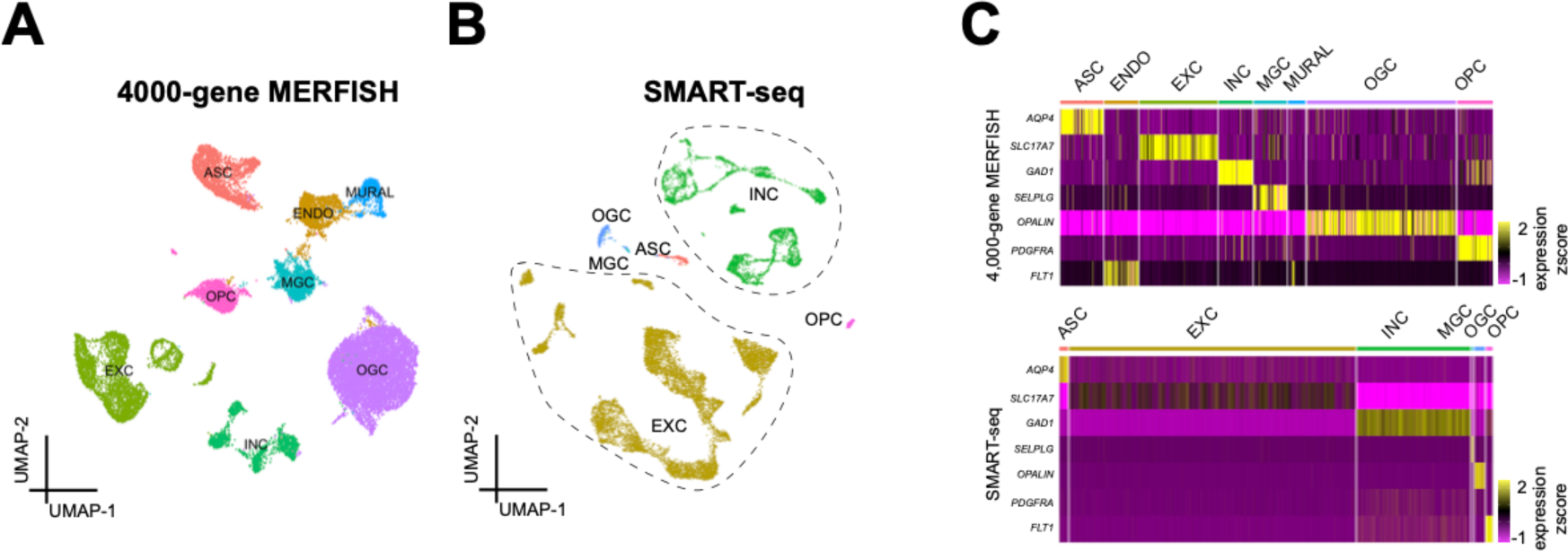
Identification of inhibitory and excitatory neurons and non-neuronal cells by MERFISH and SMART-seq. (**A-B**) UMAP visualization of inhibitory and excitatory neurons and major subclasses of non-neuronal cells identified by 4,000-gene MERFISH (A) and single nucleus SMART-seq (**B**) data. (**C**) Heatmap shows expression of marker genes for the major subclasses of cells described in (**A-B**) identified in MERFISH and SMART-seq.

**Fig. S5.**
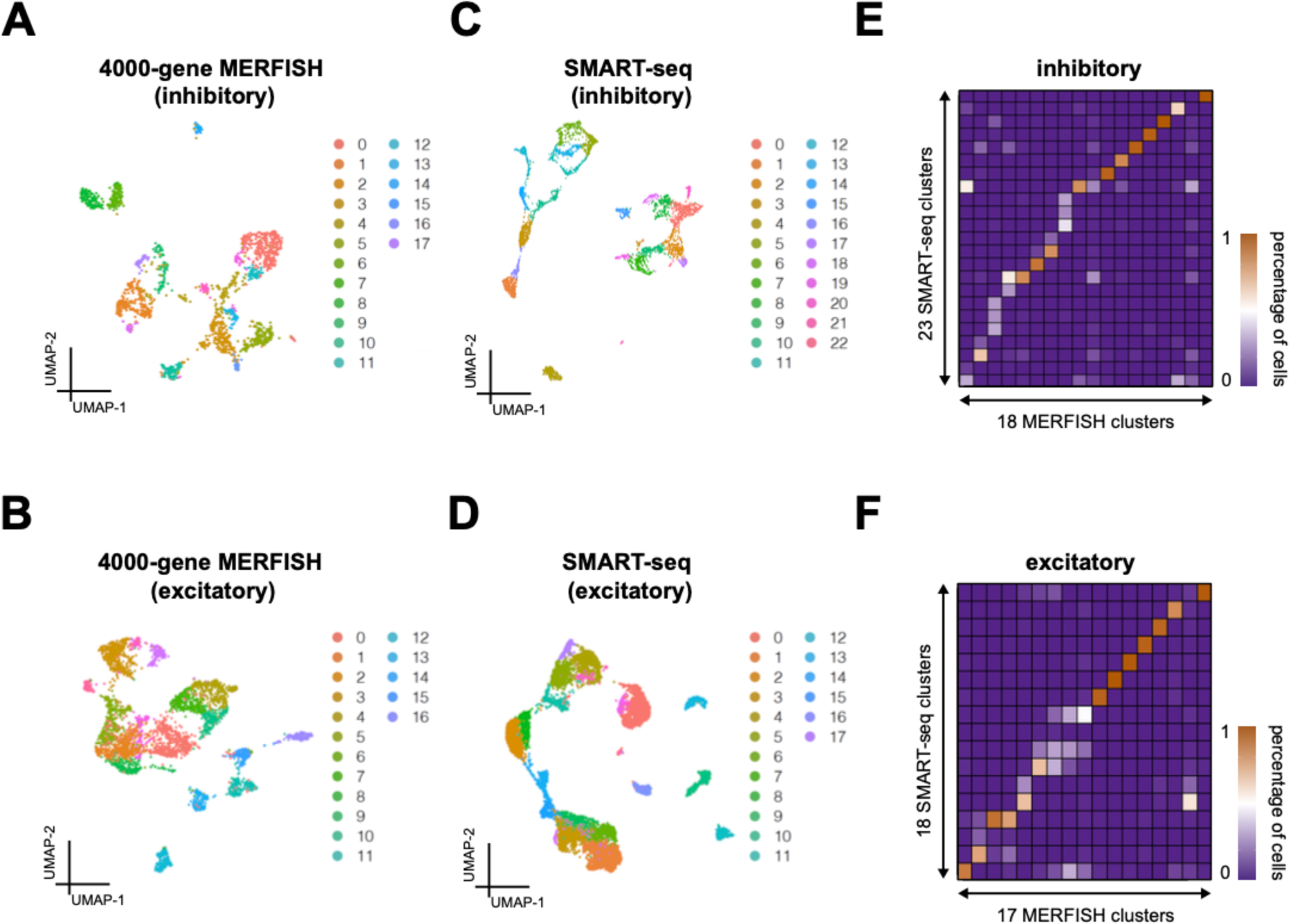
Identification of inhibitory and excitatory neuronal clusters by MERFISH and SMART-seq separately. (**A-B**) UMAP visualization of inhibitory (**A**) and excitatory (B) neuronal clusters identified by 4,000-gene MERFISH. (**C-D**) UMAP visualization of inhibitory (**C**) and excitatory (**D**) neuronal clusters identified by SMART-seq. (**E-F**) Correspondence between the clusters determined by MERFISH and the clusters determined by SMART-seq for inhibitory (**E**) and excitatory (**F**) neuronal clusters. A k-nearest neighbor classifier was trained using the MERFISH dataset and used to predict a MERFISH cluster label for each cell in the SMART-seq dataset. Cells were grouped based on their SMART-seq cluster identity, and the fraction of cells from a given SMART-seq cluster that were predicted to have each MERFISH cluster label were plotted.

**Fig. S6.**
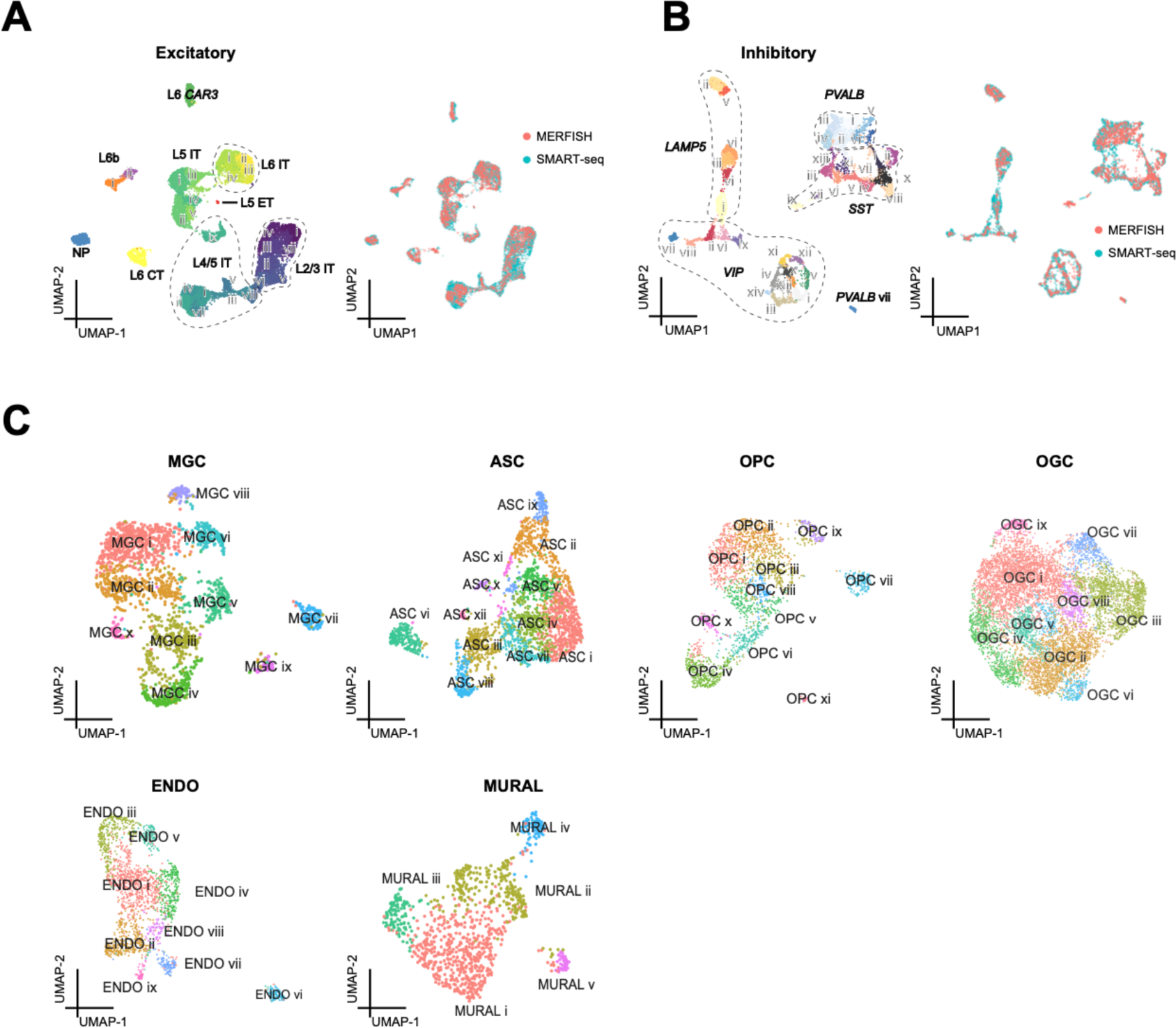
Identification of inhibitory and excitatory neuronal clusters by integrative analysis of the MERFISH and SMART-seq data and identification of non-neuronal clusters by the MERFISH data. (**A-B**) UMAP visualization of excitatory (**A**) and inhibitory (**B**) clusters identified by integrative analysis of the 4,000-gene MERFISH and SMART-seq data. Cells were colored by the cluster labels (left) or by the different technological platforms used (right). (**C**) UMAP visualization of non-neuronal clusters identified by 4,000-gene MERFISH data.

**Fig. S7.**
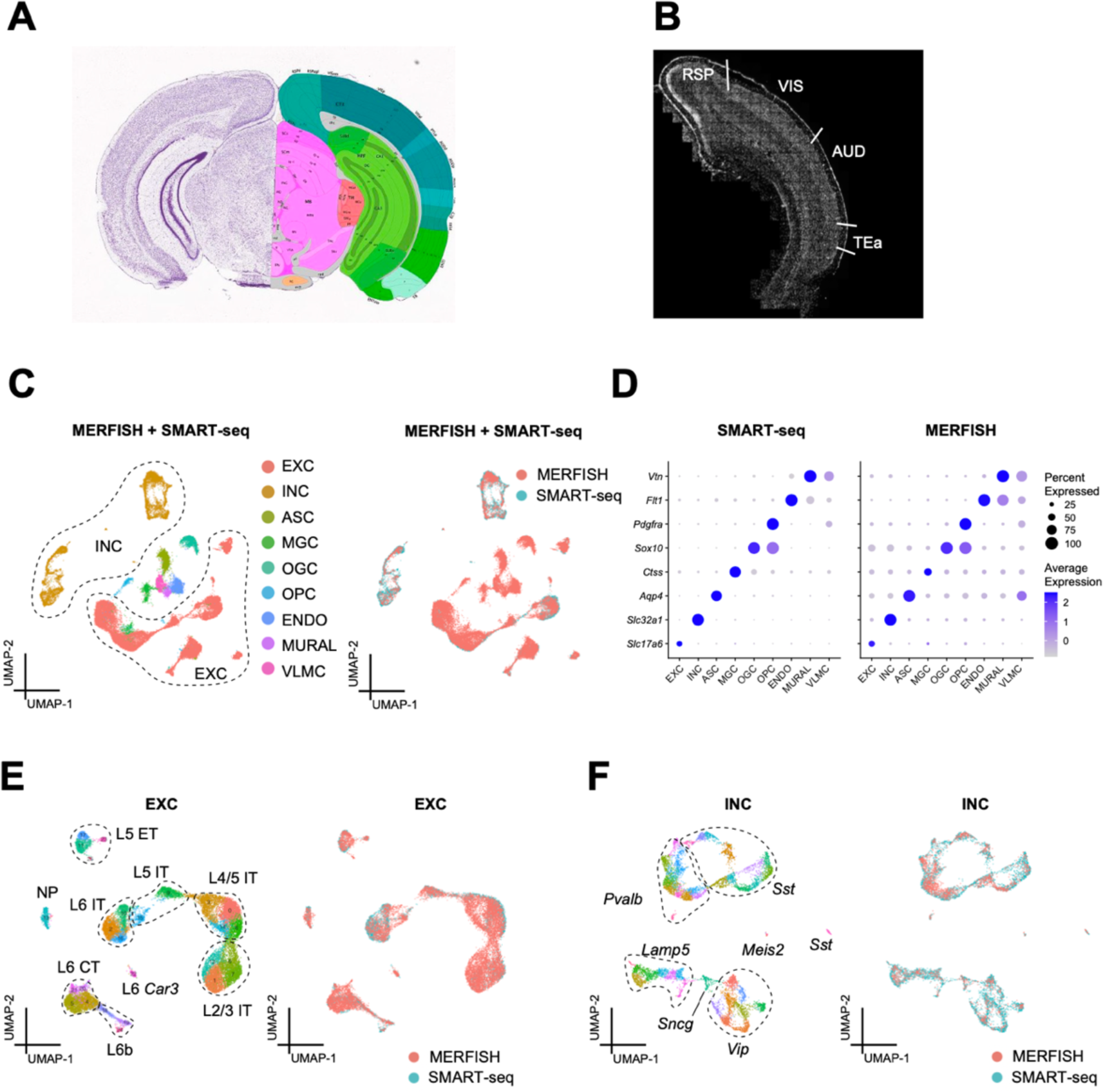
MERFISH imaging and cell type identification of the mouse cortex. (**A**) Schematic of a mouse coronal section that contains the cortical regions dissected for MERFISH imaging based on the Allen Common Coordinate Framework (CCFv2). (**B**) Mosaic DAPI image of the mouse cortical region that contained the visual cortex (VIS), auditory cortex (AUD), temporal association area (TEa), and other cortical areas imaged by MERFISH. (**C**) The inhibitory and excitatory neurons and major subclasses of non-neuronal cells identified by integrative analysis of MERFISH and SMART-seq data. Cells were colored by subclass label (right) or by technological platforms used (right). (**D**) Dot plot displaying the expression of canonical marker gene in the cell types identified in (**C**) in the SMART-seq (left) and MERFISH (right) data. The size of the dot corresponds to the percentage of cells expressing the feature in each cluster. The color indicates the average expression level. (**E-F**) The excitatory (**E**) and inhibitory (**F**) neuronal clusters identified by integrative analysis of the MERFISH and SMART-seq data. Cells were colored by cluster labels (left) or by technological platforms used (right).

**Fig. S8.**
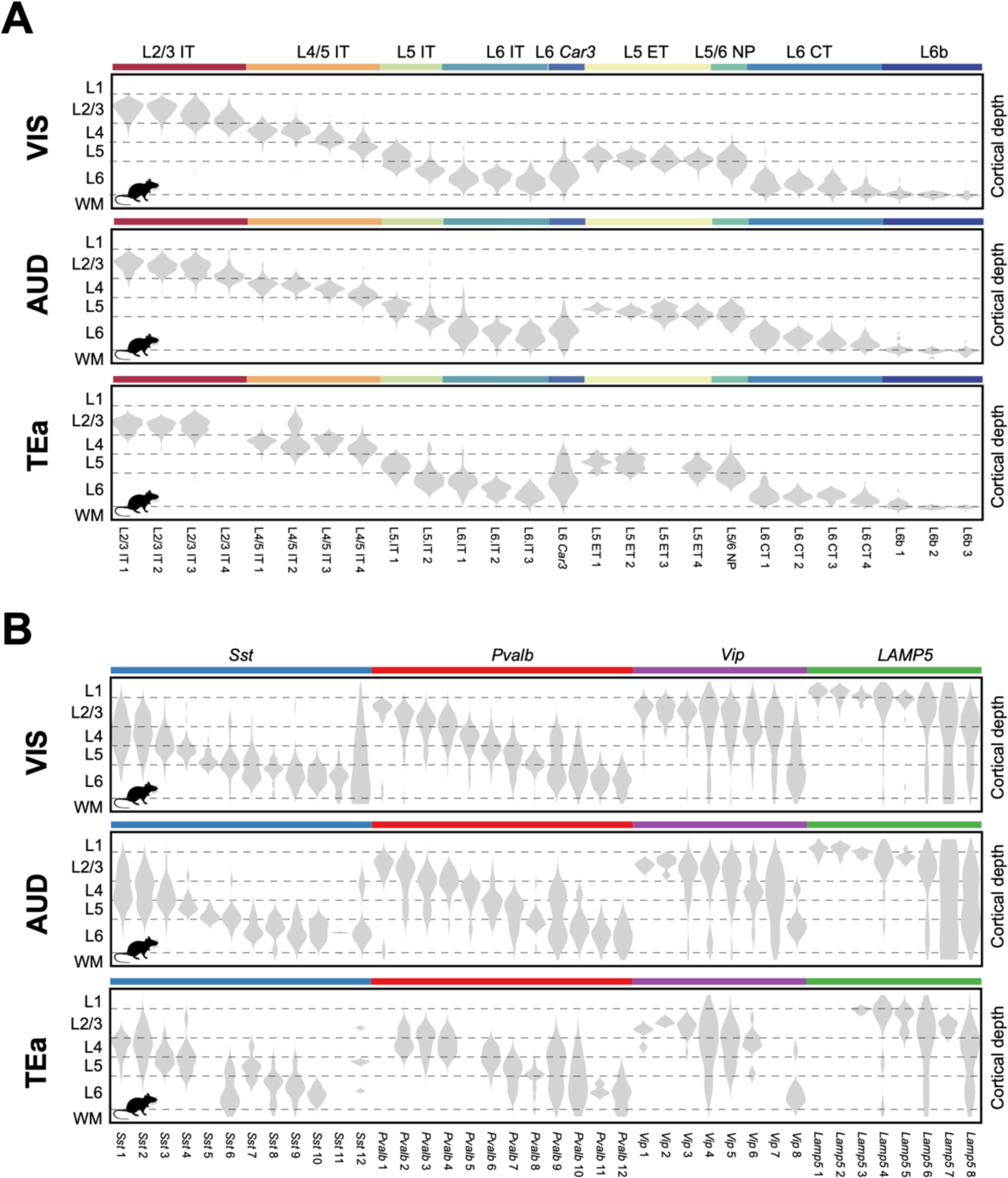
Cortical depth distributions of the neuronal clusters in different mouse cortical areas. (**A**) Excitatory neuronal clusters. (**B**) Inhibitory neuronal clusters.

**Fig. S9.**
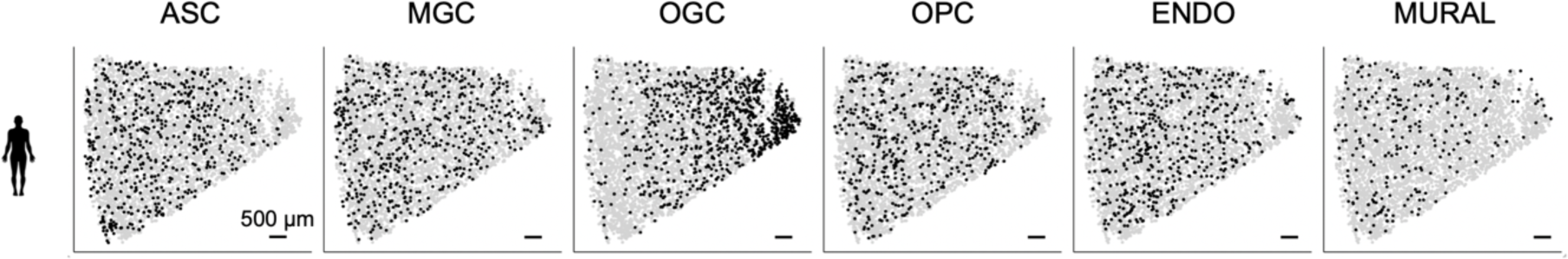
Spatial map of the non-neuronal clusters in a human MTG slice. Cells in the indicated non-neuronal subclasses are shown in black and other cells are in grey

**Fig. S10.**
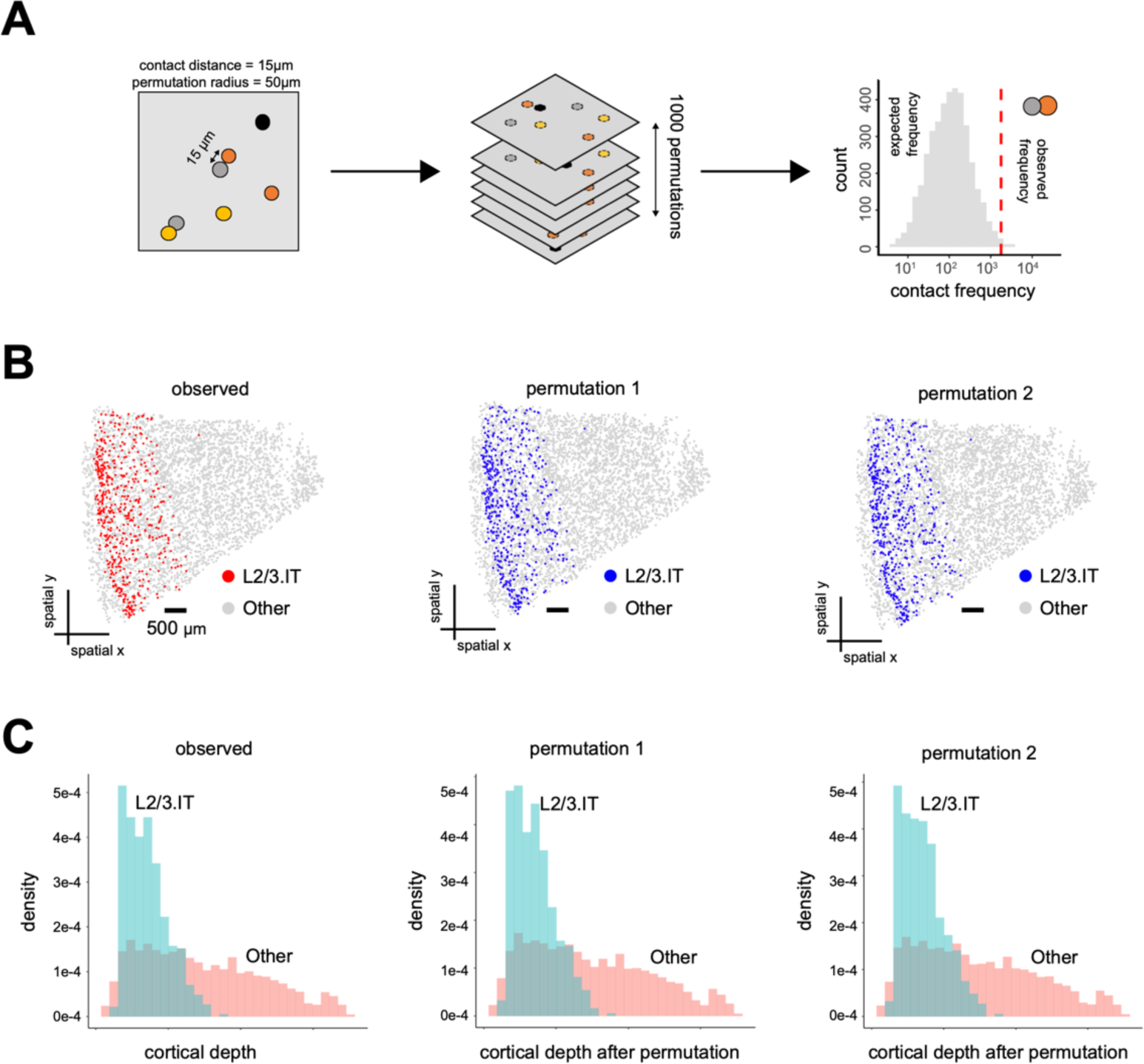
Spatial permutations that disrupt the relationship between neighboring cells while preserving the laminar distribution and local density of each cell type. (**A**) Schematic of spatial permutation test that determines the significance of interactions between cell types. Two cells were considered contacting if their nucleus centroids were within 15 μm in the imaging plane, which is approximately the size of the cell body of a single neuron. Contact frequency between any two cell types was determined as the observed frequency. Then, spatial localization of each cell was randomized within a radius of 50 μm, unless otherwise mentioned. Expected contact frequency between any two cell types was determined in each permutation and such permutation was performed 1,000 times to obtain the distribution of expected contact frequencies. The significance of observed contact frequency was calculated using one-tailed z-test and P-values were corrected to FDR (false discovery rate) using Benjamini-Hochberg Procedure. (**B**) Spatial map of L2/3 IT cells in a human MTG slice. (Left) Measured spatial map. (Middle and right) Two example spatial maps after spatial permutations described in (**A**). (**C**) Cortical depth distributions of L2/3 IT cells (light blue) and other cells (light red) in the human MTG slice shown in (**B**). (Left) Measured cortical depth distributions. (Middle and right) Cortical depth distributions after two example spatial permutations described in (**A**).

**Fig. S11.**
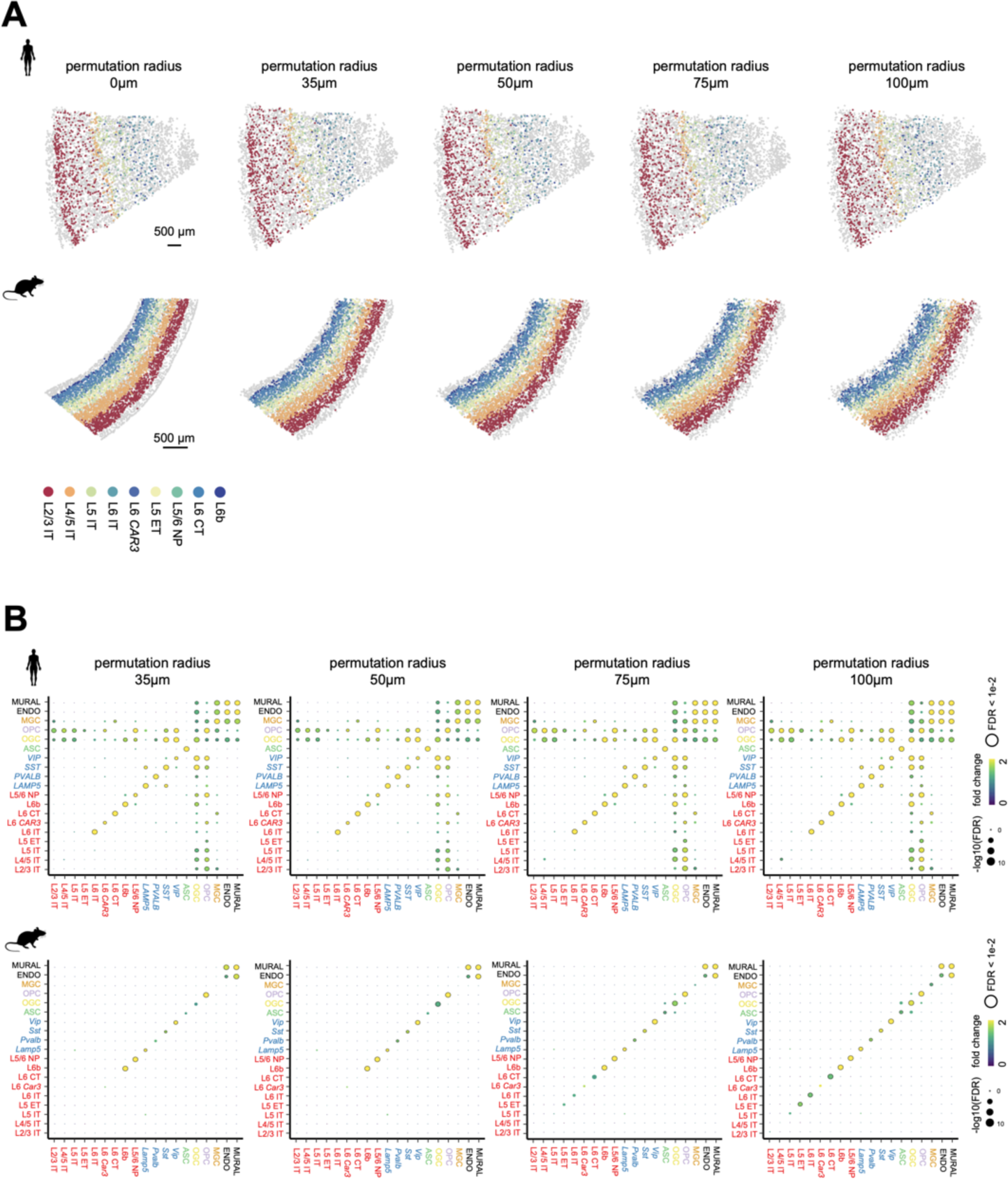
The effect of permutation radius on the spatial permutation results. (**A**) Spatial map of a human MTG slice (top) and a VIS-containing region in a mouse cortex slice (bottom) (permutation radius = 0) followed by examples of spatial maps after spatial permutations with different permutation radius ranging from 35 μm to 100 μm. Excitatory neurons were colored by their subclass labels and other cells were shown in grey. (**B**) Pairwise cell-cell interaction enrichment map for subclasses of neuronal and non-neuronal cells in human (up) and mouse (bottom) cortex shown in dot plots for different permutation radius from 35 μm to 100 μm. The map with permutation radius of 50 μm was also shown in **Fig. 3B**. The similarity between enrichment maps obtained with different permutation radii indicates the robustness of the permutation results.

**Fig. S12.**
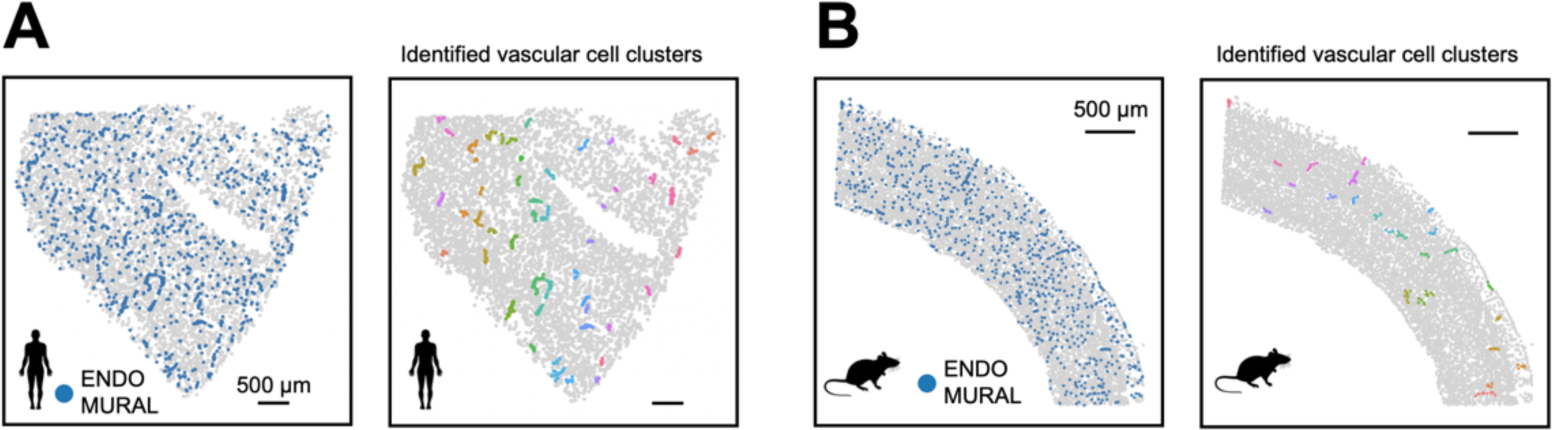
Identification of vascular structures. (**A**) (Left) Spatial map of vascular cells in a human MTG slice. (Right) Identified clusters of vascular cells in the slice. Line-shaped vascular structures, presumably blood vessels, were identified based on the density of the vascular cells and shown in colors. A k-nearest neighbor (KNN) graph was created between vascular cells in which each node represents a vascular cell, and an edge was drawn between two nodes if their distance was within 40 μm in the imaging plane. We next identified connected components in the KNN graph and components with cells fewer than six were filtered out. (**B**) As in (**A**) but for a mouse slice that contained the VIS, AUD and TEa.

## Supplementary Tables

**Table S1. MERFISH codebook for 4,000-gene measurement**. The first column is the gene name, the second column is the Ensembl transcript ID, and the following columns indicate the binary values for each of the 48 bits indicated by name of the corresponding readout sequence. Barcodes that were used as blank controls are denoted by a gene name that begins with “Blank-.”.

**Table S2. MERFISH encoding probe information for 4,000-gene measurement.** For each encoding probe, the encoding probe name and encoding probe sequence are provided. Each encoding probe name also indicates the gene name and Ensembl ID of the targeted transcript, and the names of the associated readout sequences.

**Table S3. MERFISH codebook for 250-gene measurement**. The first column is the gene name, the second column is the Ensembl transcript ID, and the following columns indicate the binary values for each of the 48 bits indicated by name of the corresponding readout sequence. Barcodes that were used as blank controls are denoted by a gene name that begins with “Blank-.”.

**Table S4. MERFISH encoding probe information for 250-gene measurement**. For each encoding probe, the encoding probe name and encoding probe sequence are provided. Each encoding probe name also indicates the gene name and Ensembl ID of the targeted transcript, and the names of the associated readout sequences.

**Table S5. Readout probe information**. For each of the bits, the bit number, the readout probe sequence name, the readout probe sequence is indicated.

